# Changes in Cerebrospinal Fluid Proteins across the Spectrum of Untreated and Treated Chronic HIV-1 Infection

**DOI:** 10.1101/2024.05.03.592451

**Authors:** Zicheng Hu, Paola Cinque, Ameet Dravid, Lars Hagberg, Aylin Yilmaz, Henrik Zetterberg, Dietmar Fuchs, Johanna Gostner, Kaj Blennow, Serena S. Spudich, Laura Kincer, Shuntai Zhou, Sarah Joseph, Ronald Swanstrom, Richard W. Price, Magnus Gisslén

## Abstract

Using the **Olink Explore 1536** platform, we measured 1,463 unique proteins in 303 cerebrospinal fluid (CSF) specimens from four clinical centers that included uninfected controls and 12 groups of people living with HIV-1 infection representing the spectrum of progressive untreated and treated chronic infection.

We present three initial analyses of these measurements: an overview of the CSF protein features of the sample; correlations of the CSF proteins with CSF HIV-1 RNA and neurofilament light chain protein (NfL) concentrations; and comparison of the CSF proteins in HIV-associated dementia (***HAD***) and neurosymptomatic CSF escape (**NSE**). These reveal a complex but coherent picture of CSF protein changes that includes highest concentrations of many proteins during CNS injury in the **HAD** and **NSE** groups and variable protein changes across the course of neuroasymptomatic systemic HIV-1 progression, including two common patterns, designated as *lymphoid* and *myeloid* patterns, related to the principal involvement of their underlying inflammatory cell lineages. Antiretroviral therapy reduced CSF protein perturbations, though not always to control levels.

The dataset of these CSF protein measurements, along with background clinical information, is posted online. Extended studies of this unique dataset will provide more detailed characterization of the dynamic impact of HIV-1 infection on the CSF proteome across the spectrum of HIV-1 infection, and further the mechanistic understanding of HIV-1-related CNS pathobiology.

## INTRODUCTION

Exposure of the central nervous system (CNS) to HIV-1 is a nearly universal facet of untreated systemic infection that is readily documented by analysis of cerebrospinal fluid (CSF) obtained by lumbar puncture (LP) [1–11]. During the earlier phases of chronic infection, CSF viral isolates are genetically similar to plasma viruses and adapted to infect CD4+ T-lymphocytes (R5 T cell-tropic viruses) [12]. As a consequence of the migration of infected CD4+ T cells from blood into the CNS, where they release virus [12–15], CSF HIV-1 RNA levels are typically maintained at levels about one-tenth of those in blood [5, 11, 16–18]. Anatomically, HIV-1 RNA detected in CSF during this period represents a sample of virus from the leptomeninges, the site of a clinically silent aseptic meningitis with mild CSF pleocytosis [19–21]. An exception to this 1:10 ratio of CSF to plasma HIV-1 RNA concentrations develops in untreated individuals with very low CD4+ T cell levels (< 50 cells per µL) in whom the ratio is closer to 1:100 when CSF white blood cell (WBC) counts are low, consistent with the concept that much of this low level CSF virus is associated with T cell trafficking [3, 4, 6, 11, 17, 22–24].

The CSF:plasma HIV-1 ratio is altered in the opposite direction in patients presenting with subacute HIV-1-associated dementia (***HAD***) in whom CSF HIV-1 RNA levels are nearly equal to those of plasma as a consequence of local viral replication within the CNS parenchyma in association with multinucleated-cell HIV-1 encephalitis (***HIVE***) [25–29]. This is generally caused by HIV-1 variants capable of infecting macrophages and related myeloid cells that express low cell-surface densities of CD4 receptors (M-tropic viruses) [30–39].

In addition to the hallmark immunosuppression that, in its severe form, underlies susceptibility to ***HAD*** and the major opportunistic complications defining the acquired immunodeficiency syndrome (AIDS), chronic HIV-1 infection is also accompanied by systemic immune activation that predisposes to an additional set of chronic morbidities [40–44]. CSF sampling in cohort studies shows that intrathecal immune activation and inflammation are also nearly constant features of chronic CNS HIV-1 infection [45]. Moreover, the character of this inflammation changes over the course of untreated infection as blood CD4+ T cells decline [45, 46]. In a recent study measuring a limited panel of inflammatory biomarkers, we identified two common patterns of CSF biomarker changes as CD4+ T cell declined during untreated infection in the absence of ***HAD***. The first pattern was characterized by a rise and then a fall in lymphocyte-related inflammation as blood CD4+ T cells decreased from >500 to below 50 cells per µL, exemplified by CSF concentration changes in CXCL10, TNF-alpha, MMP-9 and sCD14 [45]. We now refer to this ‘quadratic’ pattern of CSF viral load change with CD4+ T cell loss as the *lymphoid* pattern of biomarker change. The second ‘linear’ pattern, that we refer to as the *myeloid* pattern of CSF inflammation, was exemplified by CSF concentration changes in the macrophage inflammatory markers, CCL2 and CD163, that rose more gradually and steadily through the course of blood CD4+ T-cell decline to reach their highest levels when blood CD4+ T lymphocytes fell to <50 cells per µL in those without ***HAD***. We introduce these two terms because they serve as prototypes for similar patterns of CSF protein biomarker change noted in this analysis, as discussed later. Importantly, CSF inflammatory reactions are generally *compartmentalized* and evolve independently from the concentrations of the same biomarkers in the blood [45]. By contrast, CSF in ***HAD*** patients with the underlying multinucleated-cell form of ***HIVE*** characteristically exhibited a broad inflammatory profile with marked elevation of the CSF biomarkers that participate in both the *lymphoid* and *myeloid* CSF patterns, augmented by additional inflammatory biomarkers [45].

While CNS immune and inflammatory reactions to local infection may both aid in local viral control and facilitate viral entry, they likely also contribute to neuronal dysfunction and injury, either directly by effects on neurons or through indirect effects on uninfected myeloid cells and astrocytes by largely uncertain immunopathological pathways [47–49]. Though a number of putative mechanisms of CNS injury have been identified, the exact contributions of individual viral-encoded and inflammatory molecules to *in vivo* injury remain largely inferential [50, 51, 52., 53-62].

In addition to inflammatory changes, CSF analysis provides evidence of neuronal injury that can now be usefully monitored by measuring NfL in either CSF or blood [63–65]. Elevations of this axonal protein are not specific to ***HIVE*** [65], but in the appropriate clinical context they usefully assess ongoing *disease activity*, and complement clinical examinations and neuropsychological test documentation of cognitive-motor impairment that may be confounded by the cumulative effects of inactive (legacy) HIV-1-related and non-HIV-1-related CNS injury [66–68].

Combination antiretroviral therapy (ART) has a major impact on CNS infection, so that effective systemic viral suppression usually also induces similar CSF HIV-1 RNA suppression to below the clinical levels of detection [16, 69–71]. Indeed, even when assessed by more sensitive ‘single-copy’ assays, ART that controls plasma viremia usually suppresses CSF HIV-1 RNA to very low or undetectable levels [71, 72]. Interestingly, even in the presence of systemic treatment failure, CSF HIV-1 RNA levels usually are maintained at one-tenth or, more characteristically, even lower proportions in relation to those of plasma [17, 70]. Moreover, systemically effective ART can prevent development of HAD and usually has a salutary effect on newly-presenting HAD, curtailing clinical progression and inducing variable clinical improvement, particularly when adjusted to deliver an effective combination of drugs to the CNS site of infection [73, 74]. Monitoring reduction of CSF NfL can also be used to confirm the effectiveness of antiretroviral therapy (ART in preventing and mitigating CNS injury [68].

CSF/CNS inflammation accompanying local HIV-1 infection in the meninges and brain parenchyma is also reduced by effective ART. This includes resolution of CSF pleocytosis and reduction in the levels of various inflammatory biomarkers [17]. The dynamic effect of ART on inflammatory biomarkers has been nicely shown by rapid reduction in levels of CSF neopterin, an extensively studied inflammatory biomarker in HIV-1 infection, after treatment initiation [75, 76]. Similar overall therapeutic effects have been documented with other inflammatory biomarkers [46, 77–82].

However, CNS inflammation may not always fully subside to normal levels after suppressive treatment. For example, CSF neopterin, immunoglobulin production and T-cell activation may persist at levels above those of HIV-1-uninfected controls despite undetectable CSF HIV-1 RNA [83–86]. Even very low levels of CSF HIV-1 RNA, below the clinical cutoff of ‘undetectable’, may be associated with higher levels of local inflammatory activity than more complete CSF viral suppression [72, 76, 84, 87, 88]. Although the factors driving persistent inflammation are not firmly established, a CNS HIV-1 reservoir may be an underlying factor [15, 89–93]. Whether this persistent inflammation is harmful or protective (or both) remains to be fully defined.

There are important, more substantial exceptions to the general CSF/CNS virological treatment efficacy in the setting of systemic viral suppression. These are referred to as *CSF HIV-1 escape* syndromes in which CSF HIV-1 RNA concentrations exceed those of blood, with the levels of virus in the blood generally being suppressed by the therapy [94]. They have been classified into three distinct types: *neurosymptomatic*, *asymptomatic*, and *secondary* CSF escape (abbreviated here as ***NSE***, ***AsE*** and ***2ryE***, respectively) [94–100]. ***NSE*** is the most important of these, since it is accompanied by neurological symptoms and signs, and in some patients by severe neurological injury [95, 96, 101–106]. Its neuropathology frequently exhibits a distinct variant of ***HIVE*** with prominent CD8+ T-lymphocytes [102]. Factors contributing to the development of ***NSE*** include reduced treatment adherence, drug resistance and component drugs in the ART regime with limited CNS penetration [103–105, 107–110]. Fortunately, adjustments in ART, taking into account drug resistance, drug potency and CNS drug penetration, can usually halt disease progression and, to variable extent, reverse neurological deficits in many of these individuals [95, 96], although for some ***NSE*** patients the course can be more complicated with recrudescence [103, 105, 111]. In contrast, ***AsE***, as the name implies, is not accompanied by neurological symptoms or signs, and likely is mostly transitory, perhaps akin to viral *blips* in plasma [112–114]. It may indeed be benign, though further studies of outcomes in these individuals are needed. ***2ryE*** is provoked by other (non-HIV-1) infections within the neuraxis in which the resultant inflammatory responses activate local HIV-1 production or replication. It characteristically resolves with effective treatment of the other infection [100].

Previous CSF studies of HIV-1 infection have focused on a number of the components of local CNS inflammation in diverse cohorts and case series, often emphasizing an individual stage of infection [17, 54, 75, 115–122]. Most have also measured a relatively small number of inflammatory components [62]. For this reason, a more comprehensive picture of evolving CNS inflammation in relation to the stages of systemic infection, CNS injury, and treatment, has yet to be firmly established [22, 75, 76, 87, 123].

The aim of this study, measuring a large number of CSF proteins in parallel, was to define the features of CNS inflammation and tissue responses more broadly as they change over the course of chronic HIV-1 infection in both untreated and treated persons. To this end we used the **Olink Explore 1536** platform to measure the levels of 1463 unique proteins in more than 300 CSF samples representing the broad spectrum of clinical states defined by blood CD4+ T cells, CSF HIV-1 RNA concentrations, treatment status, and clinically recognized neurological injury during chronic infection. This report provides an introduction and initial overview of the identified CSF protein changes in relation to stages of systemic infection, local infection, and ongoing CNS injury. Importantly, the posting online of the resultant dataset opens to other investigators the opportunity to extend the value of these data by exploring additional questions related to the CSF inflammation and other protein changes that accompany the evolution of systemic infection, responses to treatment and CNS injury.

## RESULTS AND DISCUSSION

This is the initial report of an exploratory study using the **Olink Explore 1536** platform to measure a large number of CSF proteins across the spectrum of untreated and treated chronic HIV-1 infection. The resultant full study dataset is posted online for use by other investigators (**Table S1**).

The presentation of the study results and discussion is divided into two main sections. The first, ***Background*** results section, describes the clinically-defined groups in the study cohort, selected characteristics of the **Olink Explore 1536** data output illustrated by analysis of three sets of quadruplicate-repeat measurements embedded in the Olink panel, and measurements of seven CSF proteins that have been used as biomarkers of different cellular components of CNS injury in previous studies. The second, ***Main*** results section, examines salient features of the full set of measured CSF proteins in three sets of analyses that focus on: 1. overall changes in proteins across the study groups, 2. comparison of correlations of the CSF proteins with biomarkers of CNS infection and CNS injury, and 3. comparison of CSF proteins in two clinical forms of ***HIVE***, i.e., ***HAD*** and ***NSE***.

### Background Analyses

#### Study structure, study groups and their background features

This cross-sectional analysis of a convenience sample of 303 CSF samples followed a strategy used in several of our previous studies of CSF biomarkers [45, 75]. It drew on the experience and archived CSF specimens of four research centers with long-term interests in HIV-1-related CNS disease. The complete set of 307 analyzed CSF samples included 30 HIV-1-seronegative controls and 277 PLWH. The latter included 7 untreated and 5 ART-treated groups of PLWH that are the focus of this report, along with a miscellaneous group of four samples from individuals who did not clearly fit into any of the main categories and therefore were not included in this analysis which centered on group identities and differences. The background demographic and HIV-1-related clinical laboratory characteristics of the individual study participants along with the **Olink** protein measurements of the entire set of 307 CSF samples are all included in **Table S1**.

The subject group definitions are outlined in the **Methods** section, while their background characteristics are summarized in **Table S2** and graphically detailed in **Figure 1** that shows the individual values within the group designations. The format of the graphs in **Panels B** – **O** of **Figure 1** is the same as that later used to display the group-related Olink-measured CSF proteins, facilitating visual comparisons.

**Figure 1.**
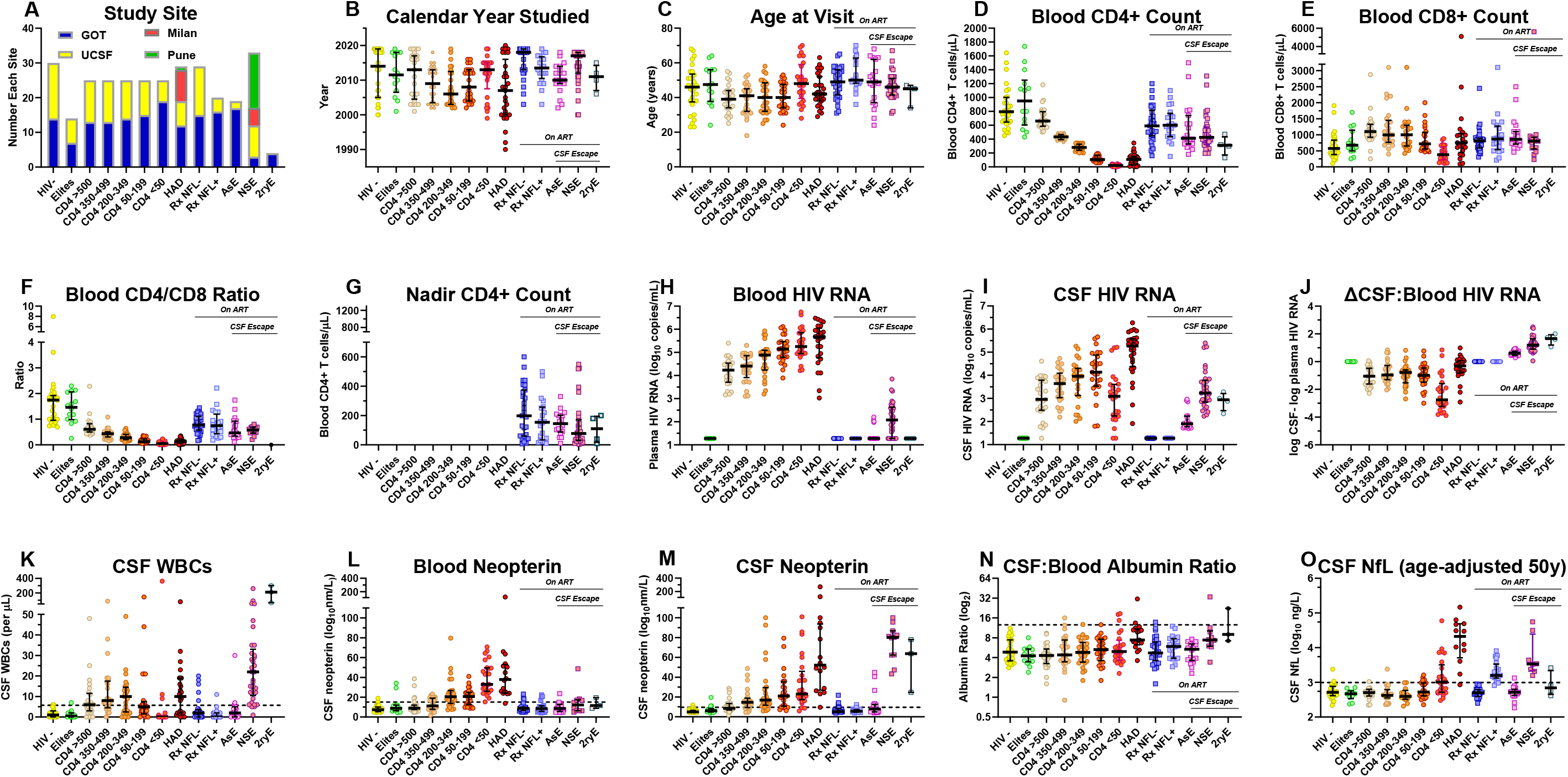
Background features of the study groups. This figure shows the details of the salient background features of the 13 study groups summarized in **Table S2**. Except for the first panel (**A**), the graphs (**B** – **O**) use the format that is applied to subsequent figures showing the Olink-generated protein measurement profiles across the study groups. This includes the symbols and the median and IQR bars. The dashed horizontal lines in Panels **K** – **O** indicate the mean +2 SD of the uninfected, seronegative (***HIV-***) control group in the measured variable as a rough estimate of their upper normal limit and as a visual guide to compare groups; this convention is also used in the Olink CSF protein measurement figures showing the group profiles in subsequent figures. **A. Study sites**. The majority of samples were obtained in the context of scheduled outpatient visits during long-term cohort studies in Gothenburg, Sweden and San Francisco, California, USA that included lumbar punctures (LPs) as part of the study protocols (Figure 1, **Panel A**). This included the specimens from the ***HIV-***, elite controllers (***elites***), five ***CD4-defined*** groups and the asymptomatic escape (***AsE***) group. The HIV-associated dementia (***HAD***), neurosymptomatic escape (***NSE***) and secondary escape (***2ryE***) samples were obtained during clinical evaluations. Additionally, the ***NSE*** and ***HAD*** groups were augmented by CSF specimens from Milan and Pune to attain a size comparable to the other main groups. When there was a larger number of CSF specimens available for a given group than required for the study, samples were chosen at random from archived collections (e.g., ***CD4-defined*** groups). An exception to this random selection was the choice of the ***HIV-*** group from Gothenburg—all were from individuals taking pre-exposure prophylaxis (PrEP) because of their self-identified risk for HIV-1 infection. ***HIV-*** controls from San Francisco were seronegative individuals from the same clinical site and demographic population as the people living with HIV (PLWH) included in the local cohort, but not on PrEP. The treatment-suppressed subjects with elevated CSF NfL concentrations (***Rx NFL+***) and the asymptomatic CSF escape (***AsE***) groups were almost all from GOT related to the local interest in these conditions. The small number of secondary escape (***2ryE***) were also all from GOT. In general, the imbalances in the subject group sizes related to scarcity of available samples (e.g., ***elites*** and ***2ryE***, resulting in smaller sample sizes), or to inclusion of larger numbers of specimens to augment particularly important groups, including the ***HIV-*** group, related to its comparative utility, and ***HAD*** and ***NSE*** groups because of our interest in more severe HIV-1-related CNS injury. These imbalances, along with those of calendar years and subject ages (**B** and **C** below), emphasize that this exploratory study used a *convenience sample* rather than one that was balanced with regard to individual demographic variables. **B.** C**alendar year**. The dates of study varied among the groups. In part this reflected the history of our cohort studies and the changes in acceptance and efficacy of treatment regimens over the collection period. These time variations impacted the length of untreated chronic infection, susceptibility to disease progression and presence of viral suppression among our study cohorts. For example, there were nearly 10 years separating the median years of the ***HAD*** group from both treatment-suppressed groups (***RxNFL-*** and ***RxNFL+***). The ***NSE*** group sampling also reflected later recognition and attention to this clinical entity. **C. Age**. The ages of the subjects were not specifically selected and consequently also varied among groups, with group medians ranging from 39 to 50 years with largely similar ranges. Overall, the median ages of the treated PLWHs were similar to those of the HIV-negative (***HIV-***) controls, while the untreated groups were generally younger. The balance of sex was not taken into account with specimen choices, so distribution also varied among the groups (**Table S2**) **D. Blood CD4+ T-lymphocytes**. The median blood CD4+ T cell count in the ***elites*** was above that of the uninfected controls and the other HIV-1-infected groups, including the two treatment suppressed groups (***RxNFL-*** and ***RxNFL+***) groups. The blood CD4+ T-cell concentrations in the 5 groups defined by these counts show the stepwise decrease from >500 to <50 CD4+ T-cells/µL dictated by the study design. The **HAD** group had a median CD4+ T cell count of 108 cells/µL that was similar to the ***50 – 199 CD4+*** group median of 106 cells/µL, in keeping with the role of advanced immunosuppression in the development of this subacute disorder and its underlying pathology of HIV-1 encephalitis (***HIVE***). The CD4+ T cell counts of the two treatment-suppressed groups (***Rx NFL-*** *and **Rx NFL +***) were nearly equal, with both having CD4+ T cell counts above those of the viremic ***CD4-defined*** groups except for those with the highest blood CD4+ T cell counts (***CD4>500*** group), indicating the partial CD4+ cell preservation or recovery with suppressive treatment. By contrast the blood CD4+ T cell counts in the three CSF escape groups were lower than the two treatment-suppressed groups, consistent with less robust recovery or ongoing loss during the escape episodes. However, these levels were not as low as in the untreated ***HAD*** group. The CD4+ blood counts of the ***2ryE*** subjects were also low, indeed below the other two escape groups, likely contributing to their vulnerability to intercurrent infections (herpes zoster in three and HSV2 meningitis in the fourth) and lead to their clinically-directed LPs. **E. Blood CD8+ T lymphocytes**. Blood CD8+ T-cell counts were on the high side of normal in the ***elites***, rose as blood CD4+ cells decreased in the untreated ***CD4-defined*** groups except for those in the ***CD4 <50*** group (median of 380 CD8+ T cells per μL). They were relatively increased in the ***HAD*** group (median 750 CD8+ T cells per μL) and in all the treated groups, including the two virally suppressed and the ***NSE*** and ***ASE*** groups (medians in 800s per μL). **F. Blood CD4+/CD8+ T cell ratio.** The differences in the trajectories of the two T cell subpopulations resulted in lowered ratios in all infected groups compared to the ***HIV-controls***, reflecting variable reductions in CD4+ and elevations of CD8+ T-lymphocytes. Reductions were most severe in the ***CD4 - defined groups***, most notably in those with CD4 counts below 200 and in the ***HAD*** group. These ratios were relatively increased in the four treated groups (0.47 to 0.77), though all remained below the level of the uninfected controls (median ratio of 1.75). Reduced ratios persisted in the two ART-suppressed groups (***RxNFL-*** and ***RxNFL+***) and the ***AsE*** and ***NSE*** groups. **G. Nadir blood CD4+ T-lymphocytes**. Nadir CD4+ T cell counts are shown only for the ART-treated groups, since CD4+ T cell values in the untreated groups at their study visits were generally at or near their nadirs. The treated groups showed varying degrees of presumed blood CD4+ T cell recovery, higher in the two virally suppressed (***Rx NFL-*** and ***NFL+***) groups (medians of 590 and 600 CD4+ T cells per μL) than in the CSF escape ***NSE*** and ***ASE*** groups (medians of 425 and 412 CD4+ T cells per μL, respectively), the latter perhaps either contributing to development of CSF escape or, alternatively, reflecting an impact of escape on CD4+ T cell dynamics. **H. CSF HIV-1 RNA**. Highest CSF HIV-1 RNA concentrations were present in the ***HAD*** group, while the ***CD4-defined*** groups showed the ‘inverted U’, or *lymphoid*, pattern of change with lower concentrations at the extremes (in the ***CD4+ >500*** and ***CD4+ <50*** groups) than in the middle ranges (***CD4+ 350***-***499***, ***200-345*** and ***50-199*** groups). We previously reported the association of this pattern with that of the CSF WBC counts as also noted in this study (see **Panel K** below), suggesting that these two findings were causally related [45]. In this study the correlation of CSF HIV-1 RNA to CSF WBC count across the ***CD4-defined*** groups was significant (Pearson correlation P = 0.006). This lymphoid pattern of infection and cell response likely underlies this same pattern in many of the CSF proteins included in the **Olink Explore 1536** panel as shown in later figures. The CSF HIV-1 RNA concentrations of three escape groups were generally below those of the untreated groups except for the ***CD4 <50*** group. CSF HIV-1 RNA concentrations were below detection in the ***elites***. **I. Blood HIV-1 RNA**. In the untreated individuals (not including the ***elites***) there was a steady increase in blood HIV-1 RNA through the full range of CD4+ T cell loss that then reached its highest mean levels in the ***HAD*** group. Unlike with CSF, there was no decrement in the ***CD4 <50*** group, showing a difference between the CSF and the systemic blood viral dynamics. The presence of low-level blood HIV-1 RNA in the ***NSE*** group may have reflected systemic partial drug resistance in some or spillover of the ‘escaped’ CNS/CSF infection into the blood. **J. CSF Blood HIV-1 RNA differences**. This panel shows the differences in viral loads between the two fluids (calculated as the log_10_ CSF HIV-1 RNA copies per mL – log_10_ plasma HIV-1 RNA copies per mL) and emphasizes the largely consistent relationship between CSF and blood HIV-1 RNA levels in untreated infection at blood CD4+ T cell levels between 50 and >500 cells/μL in which the CSF concentrations were approximately 10-fold lower than those in blood, and thus maintaining a nearly 1:10 ratio of CSF to blood HIV-1 RNA. This may relate to the kinetics in the influx of infected CD4+ T cells and of viral release into the CSF over this blood CD4+ T cell range. This ratio was disrupted when blood CD4+ T cells fell to <50 cells/μL and the CSF WBC count decreased to negligible levels; in this group the CSF:blood HIV-1 RNA ratio decreased nearly 10-fold to an overall ratio of <1:100. This underscores the importance of lymphocyte traffic and likely direct virus release by trafficking infected CD4+ T cells in the determining the CSF HIV-1 levels in the neurologically asymptomatic individuals with blood CD4+ T cell counts above 50 cells per µL. This relationship changed markedly with the development of ***HAD*** and the underlying direct neuropathic parenchymal CNS infection, i.e., ***HIVE***, in which the CSF HIV-1 RNA concentrations reached high levels, and the differences between it and the blood viral load decreased. In the ***HAD*** group local brain infection rather than hematogenous sources was responsible for the high levels of CSF HIV-1 RNA. The reversed CSF:blood HIV-1 RNA ratios in the three Escape groups were consonant with their definitions predicated on CSF > blood HIV-1 RNA concentrations that indicated the direct CNS sources of the measured CSF HIV-1 RNA in the face of systemic viral suppression. **K. CSF WBC count*s***. The median CSF cell counts rose and then fell over the course of untreated infection in the ***CD4-defined*** groups. CSF WBC counts were highest in the 200-349 group and declined to lowest levels when blood CD4+ T cells fell below 50 cells/μL, defining the *lymphoid* pattern. As discussed above, this likely was causally linked to the changes in CSF HIV-1 RNA in these groups. Both neurologically defined groups were associated with augmented local inflammation: CSF WBCs were elevated in ***HAD***, and even higher in ***NSE*** (median counts of 10 and 22 cells per μL, respectively). In these two settings the CSF WBC counts likely involved a response to local CNS infection and injury. **L. CSF neopterin**. This pteridine is predominantly, though perhaps not exclusively, a macrophage-related activation marker [151]. CSF levels showed a steady median increase as CD4 cells declined, including the highest levels in the ***CD4+ <50*** group among the ***CD4-defined*** sequence of groups, thus providing a prototypical example of the *myeloid* pattern of CSF biomarker change. CSF concentrations were even higher in the ***HAD*** group, and, notably, were highest in the ***NSE*** group, while suppressive treatment brought CSF neopterin concentrations back to near normal levels in the absence of symptomatic CSF escape. **M. Blood neopterin**. In blood, neopterin concentrations showed a similar, though more restricted elevation with CD4 decline but were notably increased in both the ***CD4 < 50*** and ***HAD*** groups, presumably indicating systemic macrophage activation in these settings that either links these two sites of infection or marks parallel changes in myeloid cell populations in the CNS and systemically. In contrast to CSF, the blood neopterin was only mildly elevated in the ***NSE*** and ***2ryE*** groups in keeping with the underlying compartmentalized CNS HIV-1 and intercurrent infections in these two groups in the face of systemic HIV-1 suppression. **N. CSF: blood albumin ratio**. This ratio provides an index of blood-brain barrier integrity, though with considerable individual and age-related variability [152]. In this sample set, only direct ***HIVE*** in the ***HAD*** group and, to a lesser degree, in the ***NSE*** group was associated with elevated median albumin ratios. Minor, though not significant, increases were present in the treated-suppressed group with elevated NFL ***(RxNFL+***). One-way ANOVA with Tukey’s multiple comparison test found significant differences (P<0.05) only for **HAD** (CD4 >500: P=0.0004; Elites: P=0.0028; AsE: P=0.0067; CD4 200-349: P=0.0069; RxNFL-: P=0.0116; CD4 50-199: P=0.0202; CD4 350-499 P= 0.0228) and **NSE** (CD4 >500: P= 0.0023); Elites: P= 0.0075; AsE: P= 0.0173; CD4 200-349 P=0.0211; HIV-1-P=0.024; RxNFL-P=0.0337; CD4 50-99 P= 0.0479). Other intergroup comparisons were not significant. Overall, the albumin ratio elevations were small, and likely to have had only limited, if any, impact on the CSF proteins measured in this panel. **O. CSF NfL.** Prior to this study, CSF NfL was measured using the UMAN ELISA method (UmanDiagnostics, Umeå, Sweden) in 209 of the 307 (71.3 percent) of the specimens. In this figure, the NfL values have been adjusted to age 50 years using the normative data and methods of Yilmaz and colleagues [68] to allow more direct subject comparison and estimations of abnormal values. While these data were incomplete and superseded in this study by the Olink measurements performed on all samples, they guided our initial definition of the ***RxNFL+*** group. The highest levels of NfL were in the ***HAD*** group, and there were elevated levels in a substantial proportion of the ***CD4 <50 group***. Notably, the NfL elevation of the ***HAD*** group had nearly a 10-fold higher median value than that of the ***NSE*** group. These observations were confirmed and extended by the measurement of NfL within the Olink Explore panel as shown later in **Figure S2**.

Despite demographic imbalances in this convenience sample (discussed in the Figure legend), the group of CSF specimens provides a robust aggregate that encompasses the main facets of chronic HIV-1 infection in the absence and presence of treatment, and importantly includes two groups with HIV-1 encephalitis (***HIVE***): the untreated HIV-associated dementia (***HAD***) and treated neurosymptomatic CSF escape (***NSE***) groups. Here we briefly focus on selected background features that particularly bear on the main CSF protein findings and on some of the interpretive themes that follow. This includes introduction to the prototypes of the *lymphoid* (CSF HIV-1 RNA and CSF WBC counts) and *myeloid* (CSF neopterin) patterns of change across the five untreated ***CD4-defined*** groups introduced earlier, along with the CSF NfL concentrations in a subset of the specimens that had been previously measured.

#### CSF HIV-1 RNA concentrations and CSF WBC counts: <u>lymphoid</u> patterns of change in the CD4-defined groups

The concentrations of HIV-1 RNA in the blood and CSF, and hence their interrelationships, changed as untreated infection progressed in the ***CD4-defined*** groups (**Figure 1**, **Panels H**, **I** and **J**). While in the blood there was a nearly steady, stepwise increase in HIV-1 RNA concentrations as the blood CD4+ T cell count decreased from the highest (***CD4 >500***) to lowest (***CD4 <50***) group levels, in the CSF progressive viral load increase was interrupted in the ***CD4 <50*** group by a drop in the median HIV-1 RNA to 3.09 log_10_ copies per mL from a peak median of 4.14 log_10_ copies per mL in the ***CD4 50 – 199*** group, resulting in a high CSF-blood difference in this group (2.75 log_10_ copies RNA per mL shown in **Panel J**). This contrasted with the more constant differences between CSF and blood HIV-1 RNA of about 1.0 log_10_ copies per mL in the other four CD4-defined groups. These changes in CSF HIV-1 RNA were parallel to, and likely related to, the CSF WBC counts (**Panel K**) in which there was also a decrease in the ***CD4 <50*** group to negligible levels in comparison to the other untreated groups with higher blood CD4+ T cell counts [17]. The low CSF WBC counts in the ***CD4 <50*** group may have related to the limited availability of both CD4+ and CD8+ T cells that generally comprise the majority of CSF WBCs in HIV-1 infection [83]. More particularly, the low number of CD4+ T cells in this group likely resulted in decreased passage of HIV-1-infected cells from the blood into the leptomeninges, and, consequently, diminished viral release into the CSF spaces. The reduced presence of T cells in this group also likely impacted the levels of some of the measured proteins in this study, resulting in this same *lymphoid* pattern for many of the CSF proteins in the sequence of ***CD4-defined*** groups, similar to the previously reported patterns in a limited subset of inflammatory biomarkers that included CXCL10, TNF-alpha, MMP-9 and sCD14 [45].

In contrast, the CSF HIV-1 RNA concentrations in ***HAD*** group increased to the highest levels among the groups, to near those in the blood (CSF median of 5.27 and blood median of 5.66 log_10_ copies RNA per mL). This was a consequence of the ***HIVE*** underlying ***HAD*** in which local CNS HIV-1 replication within the brain likely ‘spilled over’ into the CSF [17]. The median CSF WBC count in the ***HAD*** group was 10 cells per μL, and the median WBC count was even higher in the ***NSE*** group (22 cells per μL), though in the latter group the virological impact of the pleocytosis was likely partially mitigated by a treatment effect. The higher CSF than blood HIV-1 RNA concentrations in the three CSF escape groups (on therapy with virus suppressed in the blood) defined these entities, with the ***AsE*** group’s CSF virus concentrations more than 10-fold lower than the other two escape groups. Though it presents an interesting comparison to ***NSE***, the ***2ryE*** group was small and included CSF from three individuals with herpes zoster and one with HIV-2 meningitis, so it cannot be considered as representative of the larger possibilities in this category. It is also perhaps important to emphasize that CSF escape is uncommon in well-treated PLWH and that the virally suppressed groups (***RxNFL+*** and ***RxNFL-***), particularly the latter, exemplify the more typical CNS virological outcomes with current systemically suppressive ART.

#### Neopterin and the <u>myeloid</u> pattern of change in the CD4-defined groups

Neopterin is a pteridine inflammatory mediator that is likely mainly produced by myeloid and possibly to a lesser extent by astroglial cells within the CNS [124–126]. It serves as a useful CSF inflammatory biomarker during HIV-1 infection [75]. In contrast to HIV-1 RNA and WBCs, CSF neopterin showed a steady increase with falling CD4+ T cell counts in the untreated PLWH groups, without a decrease in the ***CD4 <50 group*** (**Panel L**). Based on this and previous observations with other myeloid-related biomarkers, including CCL2 and sCD163 [45], we have now termed this the *myeloid* pattern of biomarker change within the untreated groups that contrasts with the *lymphoid* pattern discussed above. We use these two convenient, albeit simplistic, terms in this report to designate recurring contrasting patterns of biomarker change in a number of the Olink-measured CSF proteins across the CD4-defined groups as presented below. The **HAD** group and two of the three escape groups, ***NSE*** and ***2ryE***, exhibited marked elevations in CSF neopterin. Blood neopterin levels also rose steadily in the untreated, though not to the high CSF concentrations found in the ***HAD*** group. By contrast, blood neopterin levels in the escape groups remained at or near normal despite the elevations in CSF, particularly in the ***NSE*** and ***2ryE*** groups, consistent with the highly compartmentalized CNS infection and inflammatory responses in these groups that contrasted with the parallel CSF and blood neopterin changes with CD4+ T cell decline in the untreated individuals [45].

#### CSF NfL in a subset of the CSF sample

**Panel O** shows the results of the CSF NfL measurements that had been assayed in a subset of the samples at the University of Gothenburg Clinical Neurochemistry Laboratory over several years in several different analytical runs of an enzyme-linked immunosorbent assay (ELISA) and age-adjusted to 50 years of age using a previously described method [68]. There were consistent NfL elevations in the ***HAD*** and ***NSE*** groups (more than ten-fold higher in the former) and milder increases in about half of the ***CD4 <50*** group, consistent with subclinical CNS injury in the setting of advanced immunosuppression. The general pattern of NfL change across this group shows similarity to the *myeloid* pattern discussed above, but a few possible differences can be noted: the elevation of NfL in the CD4 <50 group was ‘abrupt’ without a clear lead-up in the other CD4-defined groups; additionally, there was an increase in CSF NfL in the ***RxNFL+*** group that was not present with other CSF myeloid markers such as neopterin. However, similarities between the *myeloid* and this NfL, *neuronal*, pattern may also relate to the importance of myeloid cells in CNS injury.

### Olink Explore output and CSF data set

Proteins measured using the **Olink** platform are identified in this study using the gene nomenclature and numbers (**Table S1)** as they appear in the UniProt database (https://www.uniprot.org/uniprotkb/). Protein concentrations are reported in NPX units on a log_2_ scale of *relative concentrations*. Consequently, interpretations of results for the HIV-1-infected groups depended importantly on comparisons with the ***HIV-uninfected controls*** measured within the same assay. The Olink output measuring 1,472 proteins included quadruplicate measurements of three proteins (CXCL7, IL6 and TNF) as outlined below. To eliminate the effect of this redundancy, we randomly selected only one set of results for each of these three proteins to include in our analysis. This reduced the analysis to include assays of 1,463 unique proteins applied to 303 specimens, yielding a total of 443,289 individual CSF protein measurements. However, some of the measurements fell below the level of detection (LOD) of the individual assays. As noted in the Methods section, in this report we used the measured value without any correction related to the LODs in our analysis, since these low values likely had little more effect on analysis than truncating or eliminating values below the LODs. For other investigators who may want to adjust these values differently in future studies, the four LOD values for each protein assay are provided in **Table S1**.

### Analysis of three sets of quadruplicate repeat measurements

As a prelude to the main analysis, we examined the intercorrelations of quadruplicate measurements of three CSF proteins: CXCL8, IL6 and TNF. The Olink Explore 1536 platform includes four independent measurements of these proteins, one on each of the four assay plates. Within these data we also examined the impact of Quality Control (QC) Warnings that flagged some of the measurements and the frequency of protein measurements below the levels of detection (LODs) in the Olink data output file. The QC Warnings and the percentage of measurements within different strata of LODs (100; 90 - <100; 60 - <90; 40 - <60; 20 - <40; and <20 percent) are all indicated within **Table S1**.

**Figure S1A-C** show the high intercorrelations among the quadruplicate NPX values for two of the three sets of quadruplicate values and lower correlations in the third (mean R^2^ results of the 6 pairwise comparisons for each of the proteins were 0.9821 and 0.9508 for CXCL8 and IL6, with less robust correlation for the TNF correlations with a mean R^2^ = 0.7055). These correlations, along with very similar patterns of concentration changes across the subject groups (**Figure S1B**), were present despite differences in the absolute NPX values in some of the repeats, emphasizing the *relative scale* of the protein measurements and importance of a full set of embedded controls in each independent assay.

In analyzing the QC Warnings in these measurements, in all but two cases the measurements with these warnings fell within the confidence limits of the overall results. Based on this finding, and to reduce biased selection of the Olink output, we included all of the Olink results in the analyses that follow, including those with QC Warnings. Values below the LODs were minimal in the CXCL8 and IL6 repeated measures and did not impact correlations. However, more frequent results below the LODs were noted with the TNF measurements and likely contributed to less robust correlations among the four repeated measures of this protein. As noted above, for the analyses in this report, we did not exclude assay results below the limit of detection (LOD) which are indicated in **Table S1** and discussed in the legend to **Figure S1**. Values below the LODs preferentially affected the analysis of aviremic groups (e.g. ***HIV-***, ***elites***, ***Rx NFL***- and ***NFL+***), reducing the value of particular proteins to discriminate differences among these groups. Because of attention to the overall protein changes, and particularly the differences among the ***CD4-defined*** groups and the ***HAD*** and ***NSE*** groups, this issue likely had relatively limited impact on the analyses in this report. However, it is an issue that may be more important in future studies using these data when the values below the LODs will preclude more precise characterization of a number of proteins in the aviremic groups.

### Olink CNS injury markers and the use of the Olink measurement of *NEFL* as a key biomarker

In an additional preliminary study, we examined the Olink measurement of the axonal neurofilament light chain protein (NfL), that is included in the **Explore 1536** platform, along with six other biomarkers of CNS injury. In several past studies we have shown that measurement of NfL in CSF (as well as in blood) serves as a valuable biomarker of active CNS injury in HIV-1 infection [66, 68], just as it does in head injury, subarachnoid hemorrhage and a number of neurodegenerative conditions [65, 127]. Because prior CSF NfL measurements were only available from a portion of the CSF specimens included in this study (209 of the 307, 71.3 percent), we evaluated whether the Olink measurement which was available for all specimens would be suitable for use as a *background variable* in the main analyses of this study. [**NB**: In this report, we use the gene name, *NEFL*, when referring to the Olink measurement of the protein, and the more common abbreviation, *NfL*, when referring to the protein more generally or to results of measurements using other assays.]

**Figure S2**, **Panel A** shows that the profile of NEFL changes across the subject groups was very similar to that obtained with the NfL ELISA immunoassay performed in the subset of the sample shown earlier (**Figure 1**, **Panel 0**). In fact, the results of the two assays were highly correlated (R^2^ = 0.8790) (**Figure S2**, **Panel B**). These comparisons thus support the use of the NEFL results as a principal objective marker of CNS injury in subsequent analyses. We did not age-adjust the NEFL values because we had not adjusted any of the other Olink protein results and did not have sufficient data to validate age adjustment for the Olink NEFL results. The mean NEFL values in the ***RxNFL+*** group were higher than the ***RxNFL-*** group, indicating that the earlier ELISA measurements in the former group were not simply laboratory errors but reflected neuronal injury, albeit of uncertain source and meaning, particularly when not accompanied by abnormalities of other neural injury markers also shown in this figure.

The **Olink Explore 1536** platform included several other markers that have been used in previous studies to assess injury in HIV-1 infection and in other neurodegenerative settings (**Figure S2**, **Panels C - H**). Of these, CHI3L/YKL-40, TREM2 and KYNU warrant further study as useful complements to NEFL in assessing the breadth and character of CNS injury as discussed in the legend to this figure.

#### Main Olink CSF protein analyses

The following three sets of analyses explore changes in the full set of the measured CSF proteins. The first provides a broad introduction to the overall protein changes across the study groups. The second examines the relationship of these proteins to two major variables of CNS HIV-1 infection and its impact: CSF HIV-1 RNA concentration (as an index of CNS infection); and CSF NEFL concentration (as a measure of active CNS injury). The third compares two forms of ***HIVE***: ***HAD*** presenting in untreated individuals and ***NSE*** developing during treatment.

### Overview of the CSF protein changes across the spectrum of chronic HIV-1 infection

To obtain an initial, broad view of the protein changes during the course of chronic infection and begin dissecting the CSF protein changes, we performed a hierarchical cluster analysis that empirically grouped the measured proteins into three clusters, designated in **Figure 2, Panel A** as the **BLUE**, **RED** and **GREEN** clusters. The relative concentrations of the individual protein measurements are displayed vertically while the clinical groups with their individual specimens are arranged horizontally.

**Figure 2.**
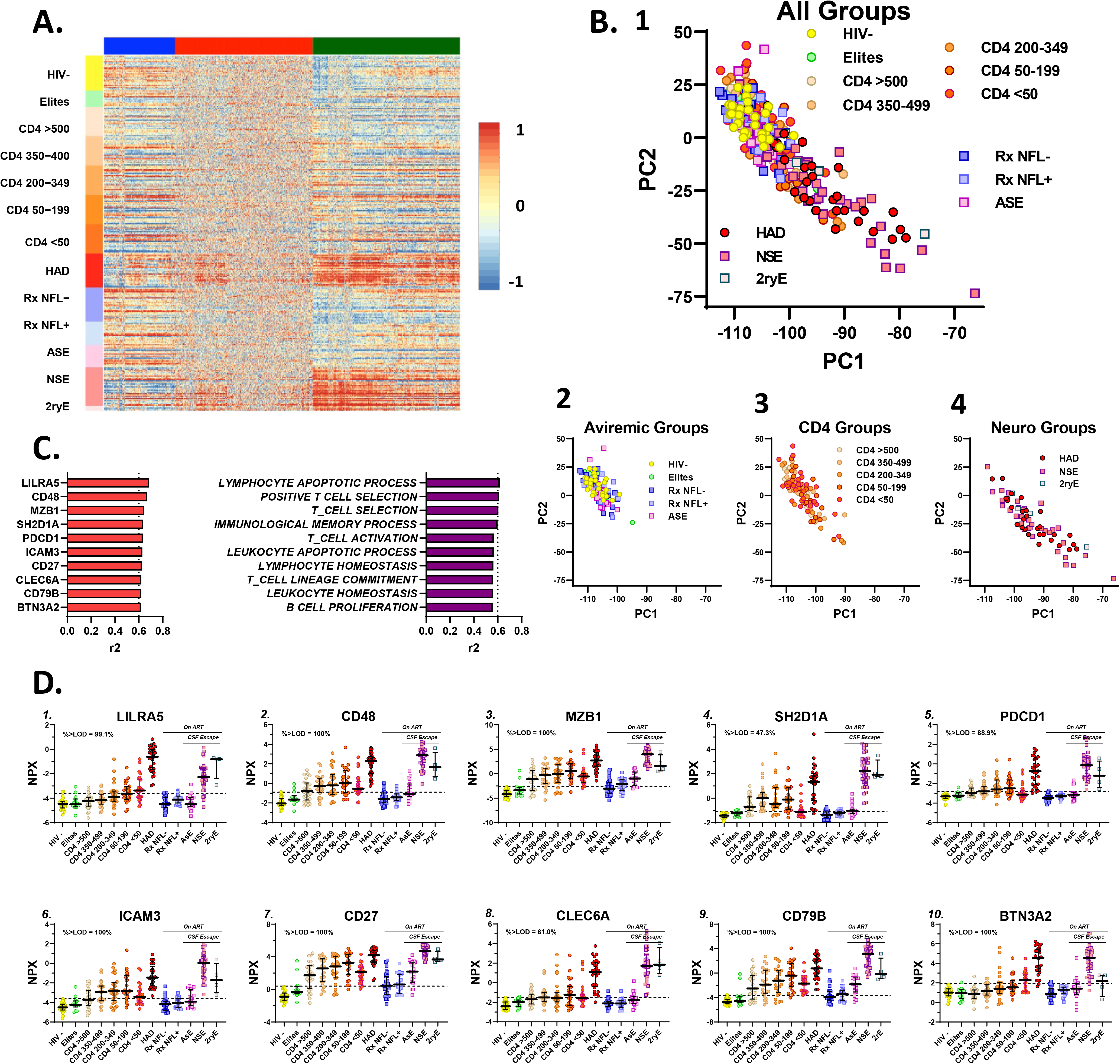
Overview of Olink Explore CSF protein measurement results. The assembled panels depict some of the main features of the aggregate CSF protein measurements using the **Olink Explore 1536** platform from different perspectives. **A.** This ***hierarchical cluster analysis*** displays the full set of the Olink CSF protein measurements in a heatmap. The study groups (and the individual CSF samples within each group) are arranged on the vertical axis, while the proteins are grouped across the figure horizontally after segregation into three clusters designated by colors at the top within **BLUE**, **RED** and **GREEN** columns of the measurements of individuals proteins in each sample. The heatmap relative scale from -1 (blue) to 0 (yellow) to 1 (red) is shown to the right of the main figure. The individual proteins were color-scaled according to their relative concentrations. The most conspicuous gradations in scale were within the **GREEN** cluster (right column) in which the many in ***HAD*** group and the two symptomatic escape groups (***NSE*** and ***2ryE***) stand out with higher (red) concentrations compared to the generally lower concentrations (blue and yellow) in the non-viremic individuals (***HIV-***, ***elites***, and ***RxNFL+*** and ***RxNFL-***) with some mixed colors over this spectrum in the ***CD4-defined*** groups. By contrast, the **RED** cluster (middle column) showed a more even and mottled spectrum except for some of the proteins shown on the left side of the cluster where there was red predominance in the ***HAD*** and two symptomatic escape groups similar to, but less pronounced than, the **GREEN** cluster. The changes among groups in the **BLUE** cluster (left column) were less distinct but with some low (blue) concentrations in the ***CD4 <50***, ***HAD*** and two symptomatic escape groups (***NSE*** and ***2ryE***), and higher concentrations in the HIV-group, i.e., showing some patterns opposite to (or inverted) in comparison to the **GREEN** cluster. Overall, the ***HAD*** and two symptomatic escape groups accounted for the greatest concentration differences from the controls among the proteins, most prominently in the **GREEN** cluster. **B.** This ***PCA analysis*** using results of all the measured CSF proteins to assess effects of the overall CSF measurements on individual subjects and their groups provides a different perspective on the protein changes across the groups. **1.** This panel includes the results from all the groups and shows a ‘gradient’ downward and to the right of the panel associated with HIV-1 systemic disease progression (as indicated by falling blood CD4+ T cell counts) and, more particularly, with the symptomatic neurological disease groups, so that the ***HAD*** and ***NSE*** individuals reach farther in this direction. Because of the superimposition of the individuals in this panel, three smaller panels (**2**, **3** and **4**) separate these groups for better definition of these changes. **2.** the aviremic groups including the ***HIV-***, the largely superimposed ***elites***, treated-suppressed (***RxNFL-*** and ***RxNFL+***) and the ***ASE*** group were all grouped together with overlap in the upper left of the plot; ***3.*** the five ***CD4-defined*** groups were also largely clustered in the same upper left area but with some individuals ‘moving’ down and to the right as immunosuppression advanced with lower blood CD4+ T-cell counts; ***4.*** the ***HAD*** and two symptomatic escape groups (***NSE*** and ***2ryE***) showed some overlap with the other groups but with further movement down and to the right (higher PC1 and lower PC2 values), again consistent with these neurological groups importantly ‘driving’ the main group differences in the aggregate CSF protein changes across the large cohort sample. Thus, these groups, defined clinically, were also, not surprisingly, separated by the changes in their CSF proteins. **C. *Influential proteins and pathways in distinguishing subject groups***. The two bar plots show the proteins and pathways that most influenced the overall differences among the subject groups. The left bar graph lists the most influential proteins, all 10 of which are markers of inflammatory processes. (Here, and elsewhere, we have followed Olink format in using UniProt abbreviations of gene designations for each protein.) The right bar graph lists the most influential defined pathways extracted from the protein changes, and again their designations emphasizes that they identify inflammatory processes, including several involving T cells. While, in part this may reflect the large number of inflammatory proteins that were included in the **Explore 1536** panel, including many with intersecting or overlapping functions, it also clearly shows that changes in inflammatory profiles determined the contours of the overall CSF proteome as systemic disease progressed and, particularly, as overt CNS disease with ***HIVE*** developed. **D. *Influential protein patterns across study subjects***. These plots show the subject group concentrations of the 10 most influential proteins (from the left side of **Panel C** above) across the subject groups using the color schema introduced in Figure 1 and used throughout this report. This includes the color symbols, medians and IQR bars, and the dashed horizontal lines indicate the mean + 2 SD of the ***HIV-* control** group in this study as an estimate of the upper limit of these variables and for visual reference across the groups. Protein concentrations are expressed in log_2_ NPX units with varying scales. The plots show the prominence of the increases in the ***HAD*** and ***NSE*** groups and their dominating influence; these two groups had the highest concentrations of all 10 of these CSF proteins, with lower and more variable patterns of elevation in the CD4-defined and other groups. As annotated in the panels, of the 10 listed proteins, the measurements in six were all (100%) above the LODs for the assay. The exceptions included LILRA5, SH2D1A, PCDC1, and CLEC6A in which 99.1%, 47.3%, 88.9% and 61.0 %, respectively, of the proteins were above the LODs (shown in the individual panels here as in subsequent figures of protein patterns across the subject groups). These influential proteins were all in the **GREEN** cluster in Figure 2A and are listed in the Glossary below with brief descriptions extracted from UniProt database (https://www.uniprot.org/uniprotkb?query<u>=*</u>) along with comments on selected features of the individual panels. **<u>Glossary of CSF proteins shown in Panel D</u>**. ***LILRA5 (A6NI73)***. (*Leukocyte immunoglobulin-like receptor subfamily A member 5*). LILRA5 may play a role in triggering innate immune responses. The ***HAD*** group had highest concentrations, but levels in the ***NSE*** and ***2ryE*** groups were also elevated above the CD4-defined groups that exhibited a gradual increase in concentrations, i.e., the *myeloid* pattern of concentration change, with CD4+ T-cell loss. ***CD48 (P09326)***. This protein is a cell surface glycoprotein that interacts via its N-terminal immunoglobulin domain with cell surface receptors, including 2B4/CD244 or CD2, to regulate immune cell function and activation; it participates in T-cell signaling. The ***NSE*** group median CD48 concentration was higher than that of the ***HAD*** group; the ***CD4 defined*** groups exhibited the *lymphoid* concentration change pattern. ***MZB1*** (***Q8WU39***). (*Marginal zone B- and B1-cell-specific protein*). MZB1 associates with immunoglobulin M (IgM) heavy and light chains, promotes IgM assembly and secretion, and helps to diversify peripheral B-cell functions. The mean concentration of the ***NSE*** group was higher than that of the ***HAD*** group. Interestingly, the levels in the ***AsE*** group were also mildly elevated, while the ***CD4+ defined*** groups showed the *lymphoid* pattern of change. ***SH2D1A*** (***O60880***): (*SH2 domain-containing protein 1A*). SH2D1A regulates receptors of the signaling lymphocytic activation molecule (SLAM) family. It also can promote CD48-, SLAMF6 -, LY9-, and SLAMF7-mediated NK cell activation. Its concentrations in the ***NSE*** group were higher than that of the ***HAD*** group and the ***CD4+ defined*** groups exhibited the *lymphoid* pattern. ***PDCD1*** (***Q15116***). (*Protocadherin alpha-C1)*. PDCD1 is an Inhibitory receptor on antigen activated T-cells that plays a critical role in induction and maintenance of immune tolerance. The concentrations in the ***NSE*** group were higher than those of the ***HAD*** group, and the ***CD4-defined*** groups exhibited a shallow *lymphoid* pattern. ***ICAM3*** (***P32942***). (*Intercellular adhesion molecule 3*). ICAM3 is in family of ligands for the leukocyte adhesion protein LFA-1 (integrin alpha-L/beta-2) and also a ligand for integrin alpha-D/beta-2. In association with integrin alpha-L/beta-2, it contributes to apoptotic neutrophil phagocytosis by macrophages. The mean concentration in the ***NSE*** group was clearly higher than that of the ***HAD*** group and the ***CD4-defined*** groups exhibited a *lymphoid* pattern. ***CD27*** (P26842). (*Receptor for CD70/CD27L*). CD27 may play a role in survival of activated T-cells and in apoptosis. The median concentration in the ***NSE*** group was slightly higher than that of the ***HAD*** group, and the CD4 groups exhibited a *lymphoid* pattern with relatively high concentrations peaking in the ***CD4 50-199*** group at levels near those of the **HAD** group. The concentrations in the ***AsE*** group were also mildly elevated, as were half of the values in the two treatment-suppressed groups indicating some inflammatory changes persisting in these groups. ***CLEC6A*** (***Q6EIG7***). (*C-type lectin domain family 6 member A*). CLEC6A **a**cts as a pattern recognition receptor of the innate immune system, drives maturation of antigen-presenting cells and shapes antigen-specific priming of T-cells toward effector T-helper 1 and T-helper 17 cell subtype. Because nearly 40% of the measurements were below their LODs, minor protein changes in the groups with lower values might have been somewhat obscured, though the overall pattern of change across groups was similar to that of PCDC1 described above with a shallow *lymphoid* pattern in the ***CD4-defined*** groups and with ***NSE*** higher than ***HAD*** group. ***CD79B*** (***P40259***). (*B-cell antigen receptor complex-associated protein beta chain*). CD79B is required for initiation of the signal transduction cascade activated by the B-cell antigen receptor complex. Here the ***CD4-defined*** groups exhibited the *lymphoid* pattern with higher concentrations in the ***NSE*** than in the ***HAD*** group. ***BTN3A2*** (***P78410***). (*Butyrophilin subfamily 3 member A2*). This protein plays a role in T-cell responses in the adaptive immune response and inhibits the release of interferon gamma from activated T-cells. The ***HAD*** and ***NSE*** group concentrations were nearly equal, with a *myeloid* pattern in ***CD4-defined*** groups.

The widest and most distinct gradations in protein concentrations were grouped in the **GREEN** cluster in which there were conspicuously high protein concentrations (at the red and orange end of the scale) in the ***HAD***, ***NSE***, and ***2ryE*** groups that contrasted with the lower concentrations (in the blue or blue mixed with yellow bands) of the aviremic ***HIV-***, ***elites***, and treatment suppressed (***RxNFL-*** and ***RxNFL+***) groups, and the more heterogeneously colored bands in the ***CD4+-defined*** groups. A similar, though less distinct and less consistent, gradation was also present on the left side of the **RED** cluster column, while the remaining two-thirds of the **RED** group showed more even concentrations (and thus narrower concentration differences) through the full range of specimens. While some of these were likely proteins that were not altered over the course of infection, others were proteins with concentrations outside of the levels of measurement of the Olink assay, mainly below the LODs (see below). The **BLUE** cluster showed more mixed gradations, included a substantial number of proteins with seemingly ‘inverted’ concentrations—i.e. with lower concentrations in the ***HAD*** and ***NSE*** groups and in the lower ***CD4 defined*** groups than in the ***HIV-controls***.

The principal component analysis (PCA) (**Figure 2B, 1-4**) provides a different perspective on the protein changes across the subject groups. It shows the impact of the overall changes in CSF proteins on separating the individual participants and their groups. The virologically undetectable PLWH (***Elites***, ***Rx NFL***- and ***NFL+***, and ***ASE*** groups), along with the ***HIV-*** control group, all largely overlapped (evident in the larger compound panel showing all the groups, **Panel A**, but more clearly in isolation in **Panel B2** showing only these groups). The ***CD4-defined*** groups also overlapped with these same groups, but with some individuals with lower CD4 counts ‘moving down and to the right’ in the compound graph and again seen more clearly in isolation (**Panel B3**). Most notably, the ***HAD***, ***NSE*** and ***2ryE*** groups ‘migrated’ still further in this same direction with greater separation of many of the samples from the other groups (**Panel B4**). Together the cluster analysis heat map and the PCA provide two consonant perspectives on the same CSF protein changes: the groups identified in the heat map as having the highest concentrations of many CSF proteins (the ***HAD***, ***NSE*** and ***2ryE*** groups) were segregated furthest from the ***HIV-*** uninfected and neurologically asymptomatic groups in the PCA analysis. Thus, these neurological conditions had a dominating effect on the CSF protein concentrations that, in turn, resulted in segregation of these neurological subject groups from the those without neurological disease.

**Panel C** in **Figure 2** lists the 10 proteins and 10 pathways that exhibited the most significant differences in CSF proteins across groups. Most of these are inflammation-related proteins, and the defined pathways involve lymphocyte-related functions as their names imply. These proteins are associated with T cell, B cell and NK cell functions. Thus, although the clinical settings of two types of neurological injury had the greatest impact on the separation of the proteins, it was the inflammatory responses in CSF in these settings that dominated their impact among the measured proteins. **Panel D** in **Figure 2** shows the group profiles of the concentration changes across the subject groups of the 10 highest correlating proteins listed in **Panel C**. While the overall patterns varied, the most conspicuous common feature was the high protein levels in the ***HAD*** and ***NSE*** groups (highest for the former in some, but in this group of proteins more frequently highest in the latter). These proteins were all from the **GREEN** group in the cluster analysis in **Panel A**, and their profiles again emphasize that the two neurological disease groups were the important drivers of the greatest CSF protein differences in the sample set. These graphs also introduce other prominent features of the CSF protein spectra across subject-group, including the two patterns of CSF biomarker change over the course of CD4+ T cell decline in the five untreated ***CD4-defined*** groups introduced earlier:

1. The *lymphoid* type, with highest levels in the ***CD4 50-199*** or ***200-350*** groups set off by lower levels in the ***CD4*** >500 and ***CD4 <50*** groups (and most notably the latter) in the ‘inverted U’ pattern discussed earlier in the context of the HIV-1 RNA and WBC concentrations. This was the most frequent pattern in this group of proteins.
2. the *myeloid type*, with progressive increases leading to highest levels in the ***CD4 <50*** group as noted earlier with CSF neopterin, was less frequent but exemplified here with the LILRA5 and BTN1A2 proteins at the upper left and lower right of this panel.

Although these ten proteins provide examples of the highest correlations, the PCA segregation of the subject group involved the entire set of measured proteins, including not only those that determined the groups separations but also those that ‘pulled’ them together because of limited protein concentrations differences. Also, while a number of these proteins shared features with the *lymphoid* profile and two with the *myeloid* profile, there were also variations within these individual profiles that bear focused and more nuanced analysis in the future.

These data illustrate the limited view provided by most previous studies, including our own, that measured only a few inflammatory mediators [2, 45, 75, 82, 128]. The results clearly underscore both the complexity and the broad participation of many proteins, including particularly many inflammatory proteins, in response to CSF/CNS infection and CNS injury. The findings, of course, do not distinguish which proteins *contribute* to CNS injury rather than simply *respond* to it. Nor do they distinguish which inflammatory proteins are involved in initiating, sustaining or determining the magnitude and character of the group inflammatory profile. These, and other, crucial mechanistic issues must be inferred and synthesized from the known functions of the proteins and their relative responses in the different clinical groups. In this report we focus on the empirical findings and leave a more fully synthesized mechanistic construction to future extensions of these analyses. Of note, at least three of these proteins showed mild elevation in the ***AsE*** group, indicating the low-level inflammatory responses in this setting that warrant future focused attention to this group.

### CSF protein relations to CNS injury (CSF NEFL) and CSF HIV-1 RNA

As a step in dissecting the forces underlying the changes in CSF proteins across the subject groups, we examined the relations of CSF proteins to two major variables that can be viewed as indicators of key *pathogenetic vectors* involved in these changes: 1. CSF HIV-1 RNA concentrations as an *index of CNS HIV-1 exposure/infection* (encompassing entry, production and replication, with the latter the main presumed drivers of CNS injury), and 2. CSF NEFL concentrations as a measure of the state of *active CNS injury* at the time of sampling. CSF HIV-1 RNA is an ambiguous indicator of CNS infection because it can reflect a variable mixture of virions derived from leptomeningeal and brain parenchymal sites of infection, depending on the clinical setting. During neuroasymptomatic infection in the ***CD4-defined*** groups (particularly those with CD4 T cell counts > 50 cells per µL), infection is largely meningeal, while ***HAD*** and ***NSE*** involve parenchymal infection (***HIVE***). However, the contributions from these sources of origin cannot be distinguished by simple measurement of HIV-1 RNA in the CSF.

Preliminary to analysis of the full set of Olink proteins, we examined the intercorrelations of CSF NEFL and HIV-1 RNA. This showed that CSF NEFL and HIV-1 RNA were, themselves, not strongly correlated across the full spectrum of study subjects (R^2^ = 0.07797) (**Figure 3, Panel A**). In part this was because the highest NEFL levels were found in the two groups with major CNS injury—the untreated ***HAD*** and treated ***NSE*** patients—that had very different levels of CSF HIV-1 RNA. Additionally, high levels of CSF HIV-1 RNA were detected in individuals in the ***CD4-defined*** groups without concomitant brain injury or elevated NEFL. Even after segregation of subsets of the groups (untreated and treated groups; and ***HAD*** and ***NSE*** groups in isolation), correlations between CSF HIV-1 RNA and NEFL remained weak.

**Figure 3.**
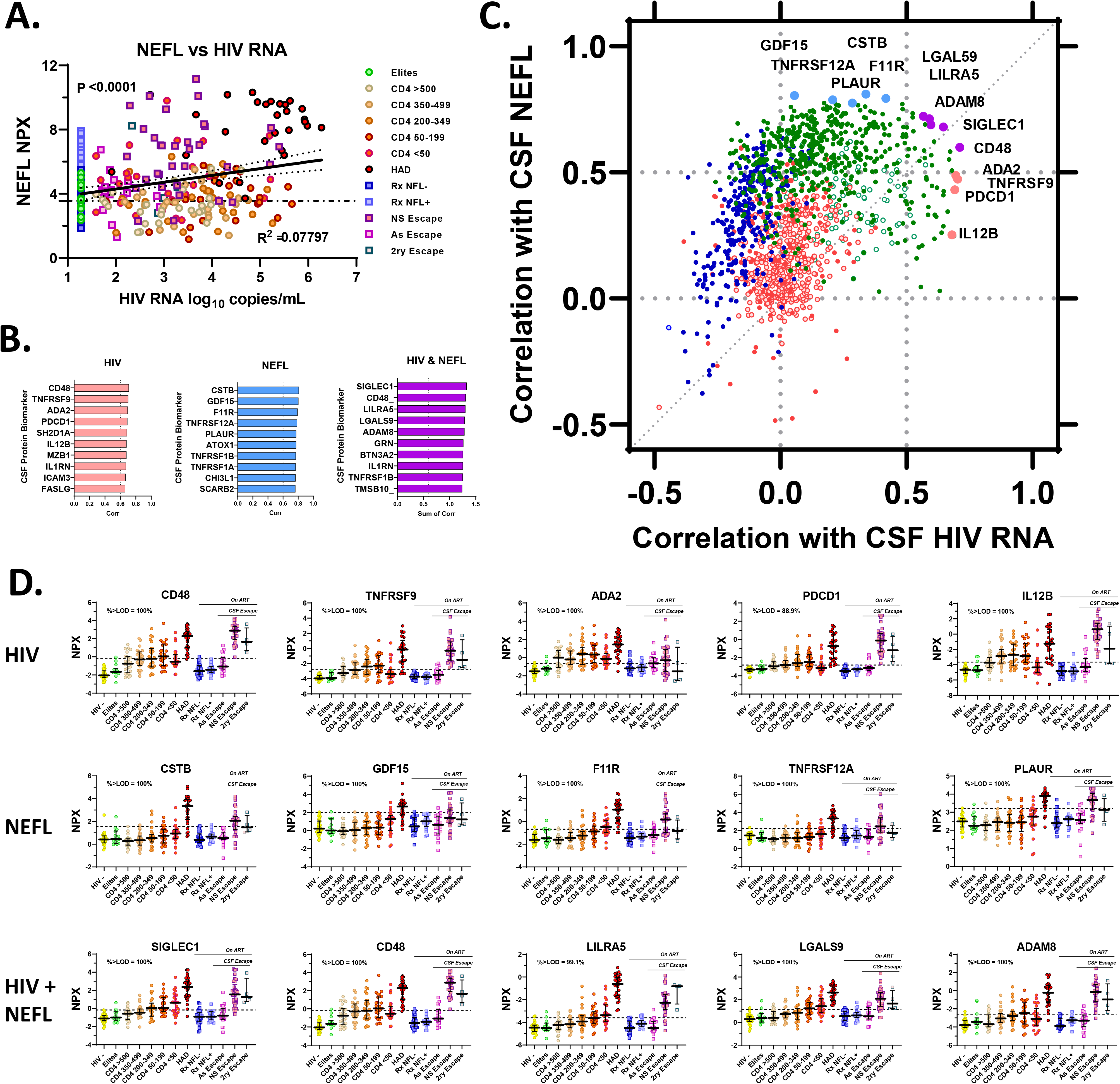
Relations of CSF proteins to CSF NfL and CSF HIV-1 RNA. This figure explores the relationships of the CSF proteins to two major variables (or *pathogenic vectors*) of HIV-1-related CNS disease: CSF HIV-1 RNA, an ambiguous index of CNS infection, and CSF NEFL (NfL measured by Olink in the ***Explore 1536*** panel), an objective measurement of active CNS injury. **A. Correlation of NEFL with CSF HIV-1 RNA**. This plot shows a significant (P<0.0001), but relatively weak correlation (R^2^=0. 0.07797) between NEFL and CSF HIV-1 RNA by linear regression. Among the features underlying this limited correlation are the high CSF RNA values without elevation of NEFL in the untreated CD4 groups, the elevations of CSF NEFL in both the treated ***NSE*** group despite their relatively lower CSF HIV-1 RNA levels (related to the partial suppression from ART) and the untreated ***HAD*** patients with high levels of HIV-1 RNA. Separate analysis (not shown) of the untreated and treated groups showed little improvement of the correlations (both also with P< 0.0001, but with R^2^ values of 0.1699 and 0.1373, respectively). Analysis of HAD patients alone showed p = 0.1709 (R^2^ = 0.07087). **B. High correlating individual proteins**. The three bar graphs show the 10 CSF proteins with: the highest correlations to ***1***. CSF HIV-1 and ***2***. CSF NEFL; and ***3***. the highest sum of the CSF HIV-1 and CSF NFL correlations. For 27 of the listed proteins, the measurements were above the LODs for the assay with some exceptions, included SH2D1A in which only 47.3% of the proteins were above the LODs and thus this protein is not included in further descriptions or in the plotting of the proteins across subject groups below. The other two exceptions were PCDC1 (88.89% above LODs) and LILRA5 (99.1 above the LODs) which are included in the further discussion and illustration. Notably some of these proteins were also previously identified as strongly impacting the overall segregation across groups as defined earlier in Figure 2. **C. Correlations of the full set of CSF proteins with CSF NEFL and HIV-1 RNA**. This figure includes a total of 1662 proteins, i.e., all of the Olink-measured proteins except NEFL, plotted according to their correlations with CSF NEFL and CSF HIV-1 RNA. It provides another broad overview of the measured proteins in the context of the two important pathogenetic vectors. Each protein symbol is colored according to their grouping in the three clusters (**GREEN**, **BLUE** and **RED**) defined earlier in the hierarchical cluster analysis (Figure 2A) with the solid symbols showing the proteins with ≥60% of the proteins above their LODs and the open symbols showing the proteins with <60% of proteins above their LODs. Exceptions to this symbol color coding are the top five proteins identified in the three groups in **Panel B** for which the symbols have been individually enlarged, color-coded using the bar colors in **Panel B** (all were originally identified in the **GREEN** cluster), and labelled with their gene names. These highly correlating proteins were located at the extremes on the right, top and right upper sections of the plot (CD48 appears in both the HIV-1 and HIV-1 + NEFL groups and is colored according to the dual correlation). The dotted grid lines mark the 0 and 0.5 correlation levels for both the Y and X axes. Overall, the **GREEN** cluster proteins populated the highest NEFL and HIV-1 RNA correlation areas with a far larger number in the ≥0.5 area of CSF NEFL correlation (Y axis) (375 proteins) than in the ≥0.5 area of CSF HIV-1 RNA correlation (X axis) (55 proteins). Twenty-two of these proteins were in the shared area in the upper right with correlations ≥0.5 for both CSF NEFL and CSF HIV-1 RNA. Some of the GREEN proteins had <60% of their measurements above the LODs as indicated by the open green circles that exhibit NEFL correlations <-.5 and CSF HIV-1 RNA correlations between 0.25 and 0.5. None of the 294 **BLUE**-clustered proteins were in the ≥0.5 HIV-1 RNA correlation area, and indeed most exhibited negative correlations with HIV-1 RNA; 52 of the **BLUE**-clustered proteins were in the ≥0.5 NEFL correlation area. Only a small number of the **BLUE**-clustered proteins showed <60% of measurements above the LODs. Of the 566 **RED**-clustered proteins 10 were in the ≥0.5 correlations with NEFL and none correlated with HIV-1 RNA ≥ 0.05. These **RED**-clustered proteins thus were concentrated along the vertical line correlation designating a 0.0 correlation with CSF HIV-1 RNA. Additionally, a large number of the **RED**-clustering proteins had <60% of proteins above the LOD (open symbols), likely contributing to their lack of correlations with either CSF NEFL or CSF HIV-1 RNA (with most grouped along the X axis at 0 correlation with CSF HIV-1 RNA). This figure with the ‘cloud’ of protein placement in relation to the two pathogenetic vectors emphasizes that the correlations of most of the Olink-measured proteins with CSF NEFL and HIV-1 RNA were largely ‘dissociated’. Only a limited number lay along the diagonal dotted line delineating equal correlations with the two major disease variables. Additionally, the much larger number of proteins correlating with NEFL at levels above 0.5 than with HIV-1 RNA, emphasizes the major influence of CNS injury on the CSF proteome. This is consistent with the strong effect of ***HAD*** and ***NSE*** on CSF proteins as can be seen by the examples below and with many of the proteins identified in other sections of the study. **D. *Patterns of high-correlating proteins across subject groups***. This Panel shows the patterns of CSF protein concentrations of the top five proteins in the three categories identified in **Panel B**. Again, the dashed horizontal lines are at the mean + 2SD of the HIV-1-group as a visual reference for comparison of group value distributions. The ***top row*** shows the five proteins with the highest correlations with HIV-1 RNA. Here the highest protein concentrations varied between the ***HAD*** and ***NSE*** groups while the ***CD4-defined groups*** exhibited a *lymphoid* pattern with a reduction in the protein levels in the ***CD4 <50*** group compared to the ***CD4 50-199*** and ***200-350*** groups. This was not surprising because this was the pattern of the CSF HIV-1 RNA concentrations and CSF WBC counts in these groups (see Figure 1**, Panels I** and **K**). The patterns in the middle row of proteins with highest correlations with NEFL exhibited two major features – highest concentrations in the ***HAD*** group and a gradual increase across the ***CD4-defined*** groups with highest values within these in the ***CD4 <50 group***, i.e., with the *myeloid* pattern. This circumstantially supports the association of CNS injury with myeloid cells. The final row with highest combined HIV-1 and NEFL correlations shows a mixture of the two patterns just outlined: variability in whether ***HAD*** or ***NSE*** had highest concentrations and in the presence of the *lymphoid* or *myeloid* patterns in the ***CD4-defined*** groups. **<u>Glossary of CSF proteins</u>**; These brief descriptions were extracted from UniProt: https://www.uniprot.org/uniprotkb/?query=*. <u>They include the top 10 proteins in each category identified in Panel B.</u> ***CD48 (P09326)***. *This cell surface glycoprotein* interacts via its N-terminal immunoglobulin domain with cell surface receptors including 2B4/CD244 or CD2 to regulate immune cell function and activation; participates in T-cell signaling. ***TNFRSF9 (Q07011)***. (*Tumor necrosis factor receptor superfamily member 9*). A receptor for TNFSF9/4-1BBL that is possibly active during T cell activation. ***ADA2 (Q9NZK5)***. (*Adenosine deaminase 2*). ADA2 **m**ay contribute to the degradation of extracellular adenosine, a signaling molecule that controls a variety of cellular responses; it may play a role in the regulation of cell proliferation and differentiation. ***PDCD1 (Q15116)***. (*Protocadherin alpha-C1*). PDCD1 is a potential calcium-dependent cell-adhesion protein. It may be involved in the establishment and maintenance of specific neuronal connections in the brain. ***SH2D1A O60880***). (*SH2 domain-containing protein 1A*). Regulates receptors of the signaling lymphocytic activation molecule (SLAM) family. Can also promote CD48-, SLAMF6 -, LY9-, and SLAMF7-mediated NK cell activation. ***IL12B (P29460)***. (*Interleukin-12 subunit beta*). Can act as a growth factor for activated T and NK cells, enhance the lytic activity of NK/lymphokine-activated killer cells, and stimulate the production of IFN-gamma by resting PBMCs; promotes production of pro-inflammatory cytokines. ***MZB1 (Q8WU39)***. (*Marginal zone B- and B1-cell-specific protein*). This protein associates with immunoglobulin M (IgM) heavy and light chains, promotes IgM assembly and secretion, and helps to diversify peripheral B-cell functions. ***IL1RN (P18510)***. (*Interleukin-1 receptor antagonist protein*). An anti-inflammatory antagonist of interleukin-1 family of proinflammatory cytokines; it protects from immune dysregulation and uncontrolled systemic inflammation triggered by IL1 for a range of innate stimulatory agents including pathogens. ***ICAM3 (P32942)***. (*Intercellular adhesion molecule 3*). A protein in the family of ligands for the leukocyte adhesion protein LFA-1 (integrin alpha-L/beta-2) and a ligand for integrin alpha-D/beta-; it contributes to apoptotic neutrophil phagocytosis by macrophages. ***FASLG (P48023)***: (*Tumor necrosis factor ligand superfamily member 6*)*. **FASLG*** binds to TNFRSF6/FAS, a receptor that transduces the apoptotic signal into cells; it is involved in cytotoxic T-cell-mediated apoptosis, natural killer cell-mediated apoptosis and in T-cell development. ***CSTB (P04080)***. (*Cystatin-B*). An **i**ntracellular thiol proteinase inhibitor. ***GDF15 (Q99988)***. (*Growth/differentiation factor 15*). A Macrophage inhibitory cytokine; regulates food intake, energy expenditure and body weight in response to metabolic and toxin-induced stresses. ***F11R (Q9Y624)***. (*Junctional adhesion molecule A*). A ligand for integrin alpha-L/beta-2 involved in memory T-cell and neutrophil transmigration; likely plays a role in epithelial tight junction formation. ***TNFRSF12A (Q9NP84)***. (*Tumor necrosis factor receptor superfamily member 12A*). A weak inducer of apoptosis in some cell types. It promotes angiogenesis and the proliferation of endothelial cells and may modulate cellular adhesion to matrix proteins. ***PLAUR (Q03405*)**. (*Urokinase plasminogen activator surface receptor*). Mediates the proteolysis-independent signal transduction activation effects of urokinase plasminogen activator. ***ATOX1 (O00244)***. (*Copper transport protein ATOX1*). May be important in cellular antioxidant defense. ***TNFRSF1B (P20333)***. (*Tumor necrosis factor receptor superfamily member 1B*). This protein mediates most of the metabolic effects of TNF-alpha. Isoform 2 blocks TNF-alpha-induced apoptosis, suggesting that it regulates TNF-alpha function by antagonizing its biological activity. ***TNFRSF1A (P19438)***. (*Tumor necrosis factor receptor superfamily member 1A*). Receptor for TNFSF2/TNF-alpha and homotrimeric TNFSF1/lymphotoxin-alpha. death-inducing signaling complex (DISC) performs caspase-8 proteolytic activation which initiates the subsequent cascade of caspases (aspartate-specific cysteine proteases) mediating apoptosis. Contributes to the induction of non-cytocidal TNF effects including anti-viral state and activation of the acid sphingomyelinase. ***CH3L1 (also YKL-40) (P36222)***. (*Chitinase-3-like protein 1*). Plays a role in T-helper cell type 2 (Th2) inflammatory response and IL-13-induced inflammation, inflammatory cell apoptosis, dendritic cell accumulation and M2 macrophage differentiation. May play a role in tissue remodeling. (See also **Figure S2**) **SCARB2 (Q14108)**. (*Lysosome membrane protein 2*). The protein acts as a lysosomal receptor for glucosylceramidase (GBA1) targeting and for enterovirus 7. **SIGLEC1 (Q9BZZ2)**. (*Sialoadhesin*). Endocytic receptor mediating clathrin-dependent endocytosis. Macrophage-restricted adhesion molecule that mediates sialic-acid dependent binding to lymphocytes, including granulocytes, monocytes, natural killer cells, B-cells and CD8 T-cells. ***LILRA5 (A6NI73).*** (*Leukocyte immunoglobulin-like receptor subfamily A member 5*). This protein may play a role in triggering innate immune responses. ***LGALS9 (O00182)***. (*Galectin-9*). Binding of this protein to HAVCR2 induces T-helper type 1 lymphocyte (Th1) death. This protein: stimulates bactericidal activity in infected macrophages by causing macrophage activation; is a ligand for P4HB and CD44; it promotes ability of mesenchymal stromal cells to suppress T-cell proliferation; expands regulatory T-cells and induces cytotoxic T-cell apoptosis following virus infection; and induces migration of dendritic cells; Inhibits natural killer (NK) cell function; enhances microglial TNF production. ***ADAM8 P78325***. (*Disintegrin and metalloproteinase domain-containing protein 8*). This protein is possibly involved in leukocyte extravasation. ***GRN (P28799)***. (*Progranulin*). GRN is a key **r**egulator of lysosomal function and a growth factor involved in inflammation, wound healing and cell proliferation; promotes epithelial cell proliferation by blocking TNF-mediated neutrophil activation preventing release of oxidants and proteases; modulates inflammation in neurons by preserving neurons survival, axonal outgrowth and neuronal integrity. ***BTN3A2 (P78410)***. (*Butyrophilin subfamily 3 member A2*). Plays a role in T-cell responses in the adaptive immune response. Inhibits the release of IFNG from activated T-cells. ***IL1RN (P18510)***. (*Interleukin-1 receptor antagonist protein*). Anti-inflammatory antagonist of interleukin-1 family of proinflammatory cytokines. Protects from immune dysregulation and uncontrolled systemic inflammation triggered by IL1 for a range of innate stimulatory agents such as pathogens. ***TMSB10 (P63313)***. (*Thymosin beta-10*). Plays an important role in the organization of the cytoskeleton.

**Figure 4.**
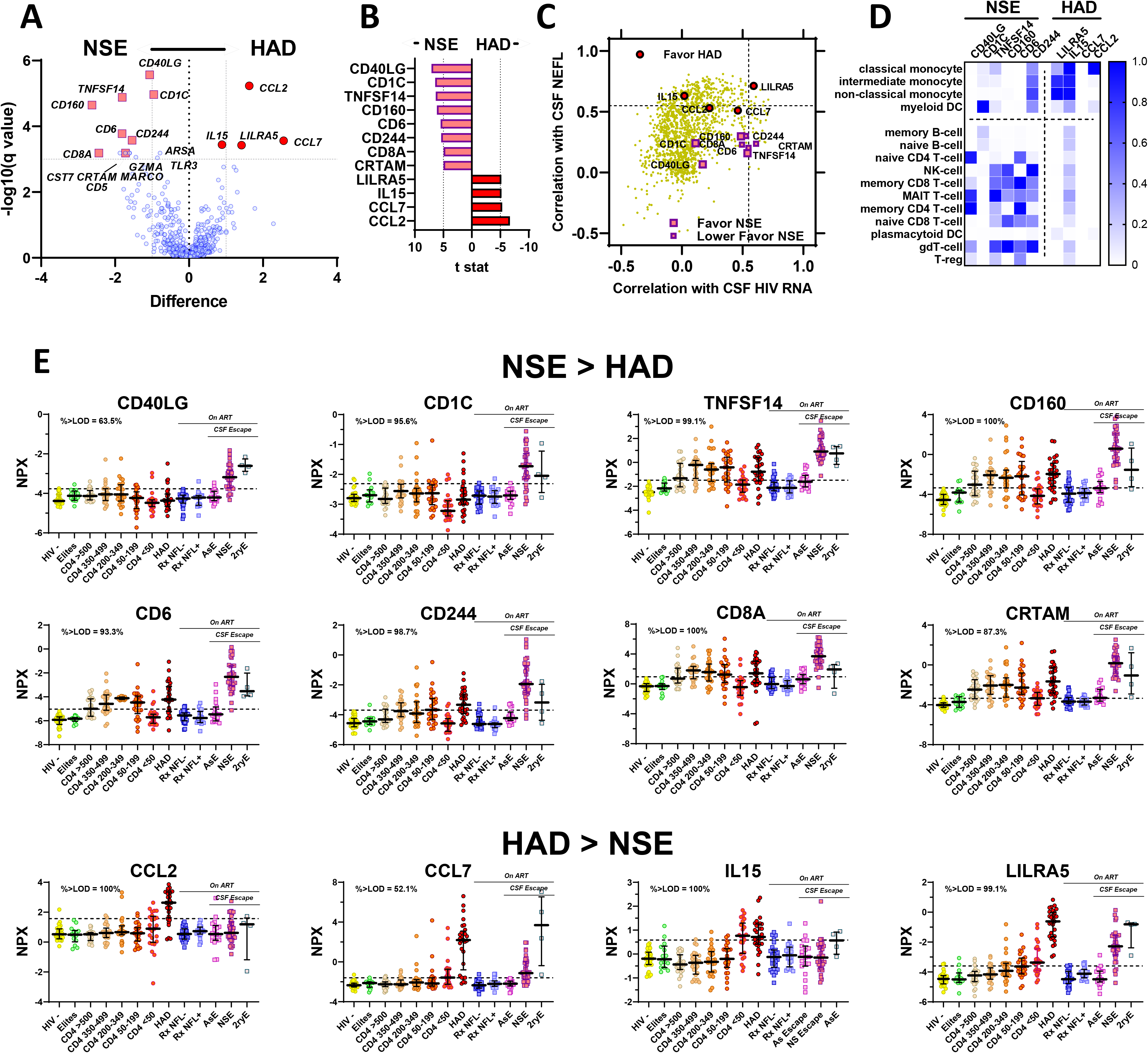
CSF proteins favoring *NSE* vs *HAD*. This figure examines CSF protein differences between ***NSE*** and ***HAD***. **A. Comparison of CSF proteins in *NSE* and *HAD groups*.** Both of these are forms of ***HIVE*** with elevations of multiple CSF proteins. For the most part, these protein elevations did not differ significantly, and this volcano plot shows a high degree of protein similarity (or lack of significant differences) within the context of the full complement of measured proteins (identified by simple blue symbols) on each side of the central vertical axis. However, there were a few identified protein differences indicated by the enlarged labelled symbols formatted using the ***NSE*** and ***HAD*** symbols that have been used in the graphs showing protein measurements across subjects. The remaining panels focus on these distinguishing proteins. **B. CSF proteins distinguishing *NSE* and *HAD* subject groups**. This panel confirms the same proteins identified by **t stat** of ≥ +/- 5 as identified in the P**anel A** volcano plot, with eight proteins favoring ***NSE*** and four favoring ***HAD***. **C. NSE and HAD distinguishing proteins in the context of their correlations with CSF NEFL and CSF HIV**. This panel shows all of the measured Olink proteins (except NEFL) in the context of their correlations with CSF NEFL and CSF HIV RNA using the same format as Figure 3C, but, for clarity, using a neutral yellow color for the full array of proteins rather than the **GREEN**, **BLUE** and **RED** designations from the hierarchical cluster analysis. The significant proteins identified in the volcano plot and t stat above (***Panels A*** and ***B***) are indicated by enlarged symbols. All were from the **GREEN** group of the cluster analysis. The proteins favoring both neurological conditions were similarly variable across a range of correlations with CSF HIV-1 RNA but were located in different vertical strata with respect to NEFL correlation. Those favoring ***HAD*** were in a range of higher range of correlation with NEFL than those favoring ***NSE***. This may relate to the generally lower levels of NEFL in ***NSE*** than ***HAD*** (Figure 2, **Panel A**). **D. Heat map of cellular associations of proteins favoring NSE and HAD.** This figure draws on literature sources describing the cell associations of 10 of the proteins identified in ***Panels A*** and ***B***, https://www.proteinatlas.org/ and [153]. The proteins listed as favoring ***NSE*** were largely associated with lymphocytes (B, T and NK lineages), while those favoring ***HAD*** associated with myeloid cells, consistent with a different balance of inflammatory cells in these two forms of ***HIVE***. **E. Different patterns of high-correlating proteins in NSE and HAD across subject groups**. The format of the graphs in this panel is the same as in previous figures showing Olink protein concentrations across the subject groups. The upper two rows plot the protein concentrations of the 8 proteins favoring ***NSE*** while the bottom row plots the proteins favoring ***HAD*** as identified in ***Panels A*** and ***B*** above. In the upper two rows, the median concentrations in the ***NSE*** group were greater than those of the comparable ***HAD*** group which were either normal or near the highest values among the ***CD4-defined*** groups. The ***CD4-defined*** groups exhibited the *lymphoid* pattern with varying levels of increase in the middle brackets. The bottom row shows the patterns of the four proteins that favored ***HAD***. Most notably, the values of ***HAD*** group were higher than those of ***NSE*** group, and the ***CD4-defined*** group patterns were either flat or had an increase in the ***CD4<50*** group above the groups with higher CD4+ T cell counts, closest to the *myeloid* or perhaps *neuronal* patterns, but with clear variation in the magnitude of differences of the ***CD4<50*** from the other ***CD4-defined*** groups. The dashed horizontal line in each panel again designates the mean +/- 2 SD of the ***HIV-*** control group for visual reference. **<u>Glossary of CSF proteins and some of their functions</u>**; Brief descriptions extracted from UniProt: https://www.uniprot.org/uniprotkb/?query<u>=*</u> ***CD40L (P29965)***. (*CD40 ligand*). Acts as a ligand to CD40/TNFRSF5; co-stimulates T-cell proliferation and cytokine production. ***CD1C (P29017)***. (*T-cell surface glycoprotein CD1c*). An antigen-presenting protein that binds self and non-self lipid and glycolipid antigens and presents them to T-cell receptors on natural killer T-cells. ***TNFSF14 (O43557)***. (*Tumor necrosis factor ligand superfamily member 14*). Acts as a ligand for TNFRSF14/HVEM to deliver costimulatory signals to T cells, leading to T cell proliferation and IFNG production. ***CD160 (O95971)***. (*CD160 antigen*). A receptor on immune cells capable to deliver stimulatory or inhibitory signals that regulate cell activation and differentiation; signaling pathways via phosphoinositol 3-kinase in activated NK cells and via LCK and CD247/CD3 zeta chain in activated T cells; receptor for both classical and non-classical MHC class I molecules; triggers NK cell cytotoxic activity, likely playing a role in anti-viral innate immune response; on CD8+ T cells, binds HLA-A2-B2M in complex with a viral peptide and provides a costimulatory signal to activated/memory T cells. ***CD6 (P30203)***. (*T-cell differentiation antigen CD6*). A cell adhesion molecule that mediates cell-cell contacts and regulates T-cell responses. It functions as costimulatory molecule and promotes T-cell activation and proliferation. ***CD244 (Q9BZW8)***. (*Natural killer cell receptor 2B4*). A receptor of the signaling lymphocytic activation molecule (SLAM) family; its ligand is CD48. It is involved in the regulation of CD8^+^ T-cell proliferation; expressed on activated T-cells. ***CD8A (P01732)***. (*T-cell surface glycoprotein CD8 alpha chain*). CD8A plays an essential role in the immune response and serves multiple functions in responses against both external and internal offenses. In T-cells, it functions primarily as a coreceptor for the MHC class I molecule:peptide complex. ***CRTAM (O95727).*** (*Cytotoxic and regulatory T-cell molecule*). CRTAM mediates heterophilic cell-cell adhesion which regulates the activation, differentiation and tissue retention of various T-cell subsets. ***CCL2 (P13500)***. (*C-C motif chemokine 2; also Monocyte chemotactic protein 1. MCP-1*). CCL2 *ex*hibits chemotactic activity for monocytes; it serves as a ligand for C-C chemokine receptor CCR. ***CCL7 (P80098).*** (*C-C motif chemokine 7*). A chemotactic factor that attracts monocytes and eosinophils. ***IL15 (P40933)***. (*Interleukin-15*). IL15 plays a major role in the development of inflammatory and protective immune responses to microbial invaders; stimulates the proliferation of natural killer cells, T-cells and B-cells and promotes the secretion of several cytokines. It induces the production of IL8 and CCL2. ***LILRA5 (A6NI73).*** (*Leukocyte immunoglobulin-like receptor subfamily A member 5*). May play a role in triggering innate immune responses.

We next examined the CSF protein correlations with the two pathogenetic vector variables, alone and in combination. **Panel B** in **Figure 3** shows the 10 proteins that separately correlated best with CSF HIV, CSF NEFL, and with both of these variables taken together (as represented by the sums of these two correlations). For a broad overview (**Figure 3**, **Panel C**), we plotted all of the Olink proteins in relation to their correlations with CSF NEFL and HIV-1 RNA, adding the color-coded grouping used in the hierarchical cluster analysis shown earlier in **Figure 2A**. Proteins from the **GREEN** cluster dominated the higher correlations with both NEFL and HIV RNA, and a higher proportion of the proteins correlated with NEFL than with HIV-1 RNA at levels above 0.5 (designated by horizontal and vertical dotted lines). This is consonant with the protein gradients noted earlier for the **GREEN** cluster showing highest concentrations in the ***HAD*** and ***NSE*** subjects and the dominant effect of these two groups on elevated CSF protein concentrations.

Some of the **GREEN** proteins had >60% of their values below the LODs, perhaps diminishing the accuracy of their correlations with either HIV-1 RNA or NEFL (though proteins in the ***HAD*** and ***NSE*** groups often exceeded the LODs even when the concentrations of the aviremic groups were below the LODs). By contrast, proteins from the **RED** cluster were mainly located in the center of **Figure 3**, **Panel C**, with HIV-1 RNA correlation values centering around zero and relatively low NEFL correlations (below 0.5) except for a few higher outliers. Many of the **RED** proteins returned >60% of measured values below their LODs, likely contributing to their frequent lack of correlation with either HIV-1 RNA or NEFL. The **BLUE** cluster was dominated by proteins with negative HIV-1 RNA correlations and NEFL correlations below the 0.5 correlation level, though with several showing negative correlations with CSF HIV RNA despite >0.5 correlation with NEFL, mingling with some of the **GREEN** cluster proteins in this location.

In **Panel C** the five highest correlating CSF proteins identified in the three bar graphs in **Panel B** are highlighted with color-coding of their locations in the plot. All of these are inflammatory proteins, and, in a broad sense, their locations emphasize the major effect of CNS injury on CSF proteins, including particularly inflammatory proteins. However, the plot also shows that HIV-1 RNA measured in CSF was associated with protein changes that had a variable relation to injury. Indeed, the highest correlations with CSF HIV-1 RNA were infrequently highly correlated with the extent of CNS injury as measured by CSF NEFL, and vice-versa, proteins correlating strongly with neuronal injury were, in general, relatively weakly correlated with HIV-1 RNA.

**Panel D** of **Figure 3** shows the group CSF profiles for the five CSF proteins with highest correlations with HIV-1 RNA, NEFL and HIV-1 + NEFL, respectively, that were identified in **Panel C**. The patterns of change in the proteins in the first two groups showed consistent differences. Those with high HIV-1 RNA correlations (top row) exhibited the *lymphoid* pattern across the ***CD4+ T cell-defined groups*** while the NEFL-correlating proteins (second row) more closely followed the *myeloid* or perhaps *neuronal* pattern of change across the ***CD4-defined*** groups with a gradual or more abrupt increase to highest levels in the **CD4 <50** group. There were variations of these patterns among these proteins; for example, PLAUR on the far right more closely resembled the NEFL’s *neuronal* pattern. In the two upper rows of graphs in **Panel D** the protein levels were more often higher in the ***HAD*** group than in the ***NSE*** group, but not in all. The combined HIV-1 and NEFL examples (third row) exhibited a mixture of the patterns seen in the other two groups (e.g., CD48 with the *lymphoid* and LILRA5 and LGALS9 with the *myeloid* patterns). Of note, the ***RxNFL+*** group did not show elevation in the inflammatory proteins that associated with NEFL, suggesting that NfL elevations in this subgroup were likely due to different processes than those involved in the two forms of ***HIVE***. This was also in agreement with the result of the other neural injury marker measurements discussed earlier and shown in ***Figure S2***.

Overall, these results emphasize that the correlations of CSF proteins with CSF NfL (in the Olink output designated as NEFL) and HIV-1 RNA were frequently *dissociated*, and that, indeed, the strongest correlations with CSF NfL were more commonly associated with relatively weak correlations with CSF HIV-1 RNA and, conversely, strong correlations with CSF HIV-1 RNA associated with relatively weaker correlation with CSF NfL. Additionally, high correlations with NfL associated with *myeloid* (and perhaps *neuronal*) patterns of change across the ***CD4-defined*** groups, while high correlations with HIV-1 RNA were associated with the *lymphoid* pattern of change across these groups. From this, we can speculate that the proteins correlating strongly with CSF HIV-1 RNA were mainly from T cells reacting to or even enhancing local HIV-1 production or replication, while the proteins correlating strongly with NfL were either effecting or reacting to CNS injury that might involve virus that evolved to grow in CNS myeloid cells, though we have not directly established these immune response-virological linkages.

### CSF proteins differences between *HAD* and *NSE*

In addition to characterizing the broad changes in the CSF proteome across the defined clinical groups, this dataset can be used to explore features that distinguish individual clinical groups or sets of groups from each other. As an initial example of this use, we compared the CSF proteins of the two groups with major HIV-1-related CNS injury: the ***HAD*** and ***NSE*** groups. Among the distinguishing background features of these two groups were: i) the absence and presence of treatment (with full or partial plasma viral suppression); ii) consequent difference in blood CD4+ T cell counts (median of 108 cells per μL in ***HAD***, and 425 cells per μL in ***NSE***); iii) difference in CSF HIV-1 RNA levels (median of 5.27 log_10_ copies per mL in ***HAD***, 3.25 log_10_ copies per mL in ***NSE***); iv) CSF WBC counts that were elevated in both but twice as high in ***NSE***); and v) different CSF NEFL concentrations (higher in ***HAD***). Previous studies also have shown that the pathology and HIV-1 genotypes/phenotypes of CSF isolates also usually differ. ***HAD*** is typically a multinucleated-cell encephalitis [129–131] involving myeloid cell infection with M-tropic viruses [132–134] developing in untreated individuals with advanced systemic infection [32, 33, 37], while ***NSE*** is generally a T-cell mediated disease with notable CD8+ T cell infiltration pathologically [102] and T-tropic HIV-1 infection (Kincer et al, unpublished observation). NSE usually occurs in the setting of limited CNS penetration of one or more antiretroviral drugs along with resistance to other drugs [94–96, 101, 103, 104, 135].

However, in common, both ***HAD*** and ***NSE*** are prominent inflammatory states within the CNS parenchyma. Both exhibited elevation of many CSF proteins. While most of these were not significantly different in the two conditions, several distinguishing proteins were identified, as shown by the volcano plot in **Figure 4**, **Panel A**. **Panel B** in this figure lists these same proteins that best distinguished the two groups with a t statistic of <5 or >5. Interestingly, those favoring ***NSE*** had a lower correlation with NEFL but, in common with the ***HAD***-favoring proteins, spanned a range of CSF HIV-1 RNA correlations (**Panel C**). Using data from the ***Human Cell Protein Atlas*** (https://pubmed.ncbi.nlm.nih.gov/31857451/), we examined reported results of studies of the expression of these proteins in purified immune cell subsets. Notably, the proteins that favored ***NSE*** appeared to relate to lymphocytic inflammation while those that favored ***HAD*** were associated with myeloid cell reactions (**Panel D**), in keeping with the differences in the pathological and virological features of these two conditions.

These differences were also reflected in the protein concentration profiles across the study groups (**Panel E**). This included the relative magnitude of the CSF protein concentrations of ***NSE*** compared to ***HAD*** and the patterns of change with CD4+ T cell loss in the five ***CD4-defined*** groups. Thus, in the top eight panels depicting the CSF proteins favoring ***NSE***, the median concentrations in ***NSE*** group exceeded that of the ***HAD*** group, with the latter ranging from normal (no different from those of the ***HIV-*** group) to levels similar to those in the highest ***CD4-defined*** group. Additionally, the ***CD4-defined*** group patterns generally exhibited the *lymphoid* pattern; in some this inverted ‘U’ pattern, though appreciable, was relatively flat, while in others it peaked at levels similar to the ***HAD*** group. By contrast, for the four proteins favoring ***HAD*** (bottom row in **Figure 4**, **Panel E**), the highest concentrations were found in the ***HAD*** group while the ***NSE*** medians were at or near normal levels in the first three shown and lower than ***HAD*** in the fourth. The values In the ***CD4-defined groups*** were either relatively flat or ascending to their highest level in ***CD4<50 group***, consistent with the *myeloid* or perhaps *neuronal* patterns. Importantly, these proteins also showed individual differences. For example, the high concentration of IL15 in the ***CD4 <50*** group, nearly equal to that of ***HAD***, was very different from the other proteins in the bottom row in which there was either an appreciable gradual, step-wise increases as blood CD4 cells decreased, or nearly flat values at CD4+ T cell counts above 50 cells per μL.

These initial comparisons were all consistent with the neuropathological differences between the two forms of ***HIVE***, with ***HAD*** typically as a myeloid-cell inflammatory and M-tropic virus-dominated process and ***NSE*** mainly a lymphocytic infiltrations and infection [102, 136], and inflammatory pathology with presumed T-tropic viruses (Kincer et al, unpublished observation). To what extent these CSF proteins simply reflect reactions to the CNS injury or, conversely, importantly participate in driving it, remains uncertain. Are the main drivers of CNS injury in the two conditions similar but the inflammatory reactions vary because of differences in the background immune system reactive capabilities, or are the mechanisms of injury more fundamentally different?

### Conclusions and future directions

This study generated an abundance of novel data related to the complex changes in CSF proteins over the course of HIV-1 infection, including after treatment. The full dataset is posted so that for other investigators can join in extending the analysis of these data. Future studies might be approached from at least two directions: 1. Continued comparisons of subject groups or subgroups to understand the distinct features of each and how they differ from selected other groups; 2. Informed analysis of the changes in particular proteins, in related proteins and in established networks and pathways to examine the mechanisms underlying the changing CSF proteome features at different phases of systemic and CNS HIV-1 infection. Inflammation is a nearly ubiquitous facet of CNS HIV-1 infection, and its relation to both systemic infection and CNS injury are clearly key aspects of CNS pathogenesis. Further definition and understanding of the CSF protein changes over the course of evolving CNS infection promises to provide enhanced understanding of the CNS pathobiology of HIV-1 infection and perhaps additional approaches to neuroprotective and neuro-mitigating therapies. It also demonstrates the potential value of applying a similar approach to other infectious, inflammatory and degenerative neurological diseases.

## MATERIALS AND METHODS

### Study design and study subjects

This was a cross-sectional, exploratory study of proteins in a convenience sample of CSF specimens obtained in research studies at four clinical centers. These specimens were derived from existing study archives and represent 11 major clinical categories, along with a small number of additional samples from uncommon clinical settings. The majority of CSF specimens were from longstanding cohort studies at the University of Gothenburg in Gothenburg, Sweden [112] and the University of California in, San Francisco California, USA [17]. These were supplemented by clinically-derived samples from ongoing HIV-1 studies in these two centers and selectively expanded by specimens from Milan, Italy [137] and Pune, India [138] representing two neurological disease groups, ***HAD*** and ***NSE***. Specimens were all obtained between 1990 and 2020 within the context of research protocols. Ethical approvals were obtained from the institutional review boards of each center (Gothenburg Ethical Committee DNr 0588-01 and 060-18; UCSF IRB Protocol 10-0727; Milan, San Raffaele Scientific Institute IRB Protocol 235/2015; and Poona hospital and Research Centre, Pune, India RECH/EC/2018/19/257). The samples from the cohort participants and the patients in the disease groups were obtained after written informed consent; if patients were unable to directly consent (some of those with ***HAD*** and ***NSE***), this was obtained from a person with power of attorney or the equivalent. The sample groups included an uninfected control group and 12 chronically HIV-1-infected groups defined by blood CD4+ T-cell counts, neurological presentations, treatment status and viral suppression as previously described [123]. When samples exceeded the planned number within each of these predefined groups, specimens were chosen at random, while in groups with sparse samples all available specimens were included. In brief, the following were the group criteria (the final specific individual group characteristics are described in the **RESULTS AND DISCUSSION** section).

#### HIV-1 uninfected controls (*HIV-*)

were drawn from the Gothenburg and San Francisco populations with similar demographic and background characteristics to the PLWH at those centers. Notably, the Gothenburg specimens were selected from a group of individuals taking pre-exposure prophylaxis (PrEP) because of HIV-1 infection risk. As with the PLWH, all of these participants volunteered for the clinical examinations, background blood tests and the study lumbar punctures (LPs). All were screened with HIV-1 serology and blood HIV-1 RNA measurements.

#### HIV-1-infected, untreated

included 7 groups of PLWH.

##### Elite viral controllers (elites)

were defined by positive HIV-1 serology and undetectable plasma HIV-1 RNA concentrations on repeated occasions using a standard clinical cutoff (<20 HIV-1 RNA copies per mL) in the absence of ART [23, 24, 139].

##### Five CD4-defined groups

were PLWH who did not have evidence (symptoms or signs) of active or confounding neurological disease on clinical examination. They were segregated by their blood CD4+ T lymphocyte count into five strata: **>500**, **350-499**, **200-349**, **50-199**, and **<50** blood CD4+ T-lymphocytes per µL, respectively.

##### HAD

patients were PLWH who presented clinically with new symptomatic, subacute neurological disease attributed to CNS HIV-1 infection by their caring physicians based on their clinical presentation and diagnostic evaluations. Some were studied before publication of the formal Frascati criteria for ***HAD*** [140] and diagnosed with AIDS dementia complex (ADC) stages 2-4 [131] while also meeting the American Academy of Neurology criteria for HIV-1-related dementia in place at the time [141]. Retrospectively, they also met the functional criteria for the Frascati diagnosis of ***HAD*** without the requisite formal neuropsychological assessment, and therefore this term was used to encompass the full group of these patients.

##### HIV-1-infected, treated groups

included 5 main groups of PLWH taking ART.

Two groups of treated, *virally suppressed* individuals were selected from the Gothenburg and San Francisco cohorts: one with either normal age-corrected CSF NfL or previously unmeasured CSF NfL (***RxNFL-***), and a second group of special interest with elevated age-corrected CSF NfL (***RxNFL+***) levels on previous testing.

##### CSF escape

was diagnosed in individuals taking ART with suppression of plasma HIV-1 RNA below pretreatment levels but with higher concentrations of HIV-1 RNA in CSF than plasma [94, 113, 135]. Three types of CSF escape were distinguished: ***NSE*** in which individuals presented clinically with new neurological symptoms and signs that were attributed to CNS HIV-1 infection and without alternative cause after diagnostic studies; asymptomatic escape (***AsE***) were neurologically asymptomatic individuals identified by CSF and blood HIV-1 RNA findings during participation in cohort studies; and secondary CSF escape (***2ryE***) was diagnosed on the basis of CSF and plasma measurements in the context of another, non-HIV-1 nervous system infection [100].

##### Miscellaneous PLWH

included four anecdotally interesting individuals who did not fit the above categories: one who sustained HIV-1 cure (the ‘Berlin patient’) [142], two with very early initiation of ART [143], and one ambiguous escape with a history of PML. These individuals are included in the full data table (**Figure S1)** but not in the analyses described in this report.

##### CSF Sampling

CSF from the cohort subjects was obtained according to previously described standard protocols [17, 112, 144, 145]. Similar methods were used to process clinically-obtained samples from **HAD**, **NSE** and **2ryE** groups that were preserved for study purposes. In brief, CSF, obtained by LP, was placed immediately on wet ice and subjected to low-speed centrifugation to remove cells before cell-free fluid was aliquoted and stored within 2 hours of collection. Samples were maintained at ≤-70°C until the time of HIV-1 RNA and biomarker assays.

### Clinical Evaluations and Background Laboratory Methods

Cohort subjects underwent standardized general medical and neurological assessments at study visits as previously described [146]. Symptomatic CNS disorders (***HAD***, ***NSE***, and ***2ryE***) were evaluated using standard clinical and neurological assessments, including neuroimaging along with CSF and blood measurements, to confirm their diagnoses and assess disease severity [146]. All cohort subjects had routine clinical bedside screening for symptoms or signs of abnormalities in cognitive or motor function or evidence of CNS opportunistic infections or other conditions that might impact CSF biomarker concentrations. Individuals with CNS opportunistic infections or other conditions confounding these analyses were omitted with the exception of those in the ***2ryE*** group in which these infections were part of the clinical group definition. We initially set a general target of 25 separate samples for each major clinical group. Subsequently, some groups of particular interest were augmented with additional specimens, while for other categories there were shortfalls related to limited specimen availability.

HIV-1 RNA levels were measured in cell-free CSF and plasma at each site using the ultrasensitive Amplicor HIV-1 Monitor assay (versions 1.0 and 1.5; Roche Molecular Diagnostic Systems, Branchburg, NJ), Cobas TaqMan RealTime HIV-1 (version 1 or 2; Hoffmann-La Roche, Basel, Switzerland) or the Abbott RealTime HIV-1 assay (Abbot Laboratories, Abbot Park, IL, USA), depending on the time period of collection and the local Clinical Research Laboratories. All recorded viral loads reported below 20 copies per mL were standardized to an assigned ‘floor’ value of 19 copies per mL for descriptive and computational purposes. Each cohort study visit included assessments by local clinical laboratories using routine methods to measure CSF white blood cell counts (WBCs), blood CD4+ and (in many) CD8+ T lymphocyte counts by flow cytometry, and CSF and blood albumin to assess blood-brain barrier integrity. CSF neurofilament light chain protein (NfL) concentrations had been measured in multiple runs in a proportion of the subjects prior to this study using a sensitive sandwich enzyme-linked immunosorbent assay (NF-light^®^ ELISA kit, UmanDiagnostics AB, Umeå, Sweden) [147] performed in the Clinical Neurochemistry Laboratory at the University of Gothenburg by board-certified laboratory technicians blind to clinical data; intra-assay coefficients of variation were below 10% for all analyses. To compare these NfL values across all groups, we calculated age-adjusted NfL values for 50 years of age and compared them to the 50 year-old upper limit of normal of 991 ng/L, as outlined by Yilmaz and colleagues [68].

### Olink Explore methods

Samples from all sites were maintained at ≤-70° C, aggregated in San Francisco and shipped together to Olink Laboratory in Watertown, MA, USA in previously stored, frozen aliquoted tubes, and analyzed in November 2020. Samples were transferred using local Olink procedures. The ***Explore 1536*** battery deployed at that time consisted of four plates: *Inflammation, Cardiometabolic*, *Oncology* and *Neurology* plates. The Olink immunoassays are based on the Proximity Extension Assay (PEA) technology [148], which uses a pair of oligonucleotide-labelled antibodies to bind to their respective target protein. When the two antibodies are in close proximity after binding, a new polymerase chain reaction target sequence is formed, which is then detected and quantified by Next Generations Sequencing (NGS). Output is reported in NPX log_2_ units which are used for within-assay comparison of protein concentrations (https://olink.com/).

The level of detection (LOD) for each assay was listed in the data output and was based on the background, estimated from the negative controls on every plate, plus three standard deviations. In the results presented below for this analysis, we included the returned values without adjustment or substitution when below the LOD. This is one of three strategies discussed by Olink and adopted for this study both for simplicity, and because it may increase the statistical power and provide a more normal distribution of the data without an increase in false positives (https://olink.com/faq/how-is-the-limit-of-detection-lod-estimated-and-handled<u>/</u>). Each protein measurement was comprised of four subgroups with a related LODs; these are listed in **Table S1**. We also included measurements that were flagged in the data output file with quality control (QC) Warnings. These Warnings were generated if incubation Control 2 or Detection Control (corresponding to that specific sample) deviated by more than a pre-determined value (+/- 0.3) from the median value of all samples on the plate (https://olink.com/faq/what-does-it-mean-if-samples-are-flagged-in-the-qc/). Our own analysis of the effect of the QC warnings in the three proteins with quadruplicate repeats supports inclusion of these values is shown and discussed in the context of **Figure S1** in the **Results**/**Discussion** section.

### Statistics analysis

R version 4.1.1 was used for the main data analysis. The “hclust” function in R was used to perform hierarchical clustering to identify three clusters of the proteins. The “prcomp” function in R was used to perform the principal component analysis. One-way ANOVA was used to test the differential expression of proteins between the clinical groups. Biomarker associations were analyzed across the entire sample set using Spearman’s rank correlation, graphically presented as a heat map for some. Comparison of biomarker concentrations between subject groups to address selected *a priori* questions used non-parametric methods including Mann-Whitney to compare two groups and Kruskal-Wallis with Dunn’s *post hoc* test to compare three or more groups.

Two-sample t tests were used to compare the protein expression between the ***HAD*** and ***NSE*** groups. Partial correlation was used to assess the association between protein markers and the CSF HIV-1 level while controlling for the CSF NfL level (*NEFL* in Olink output). Similarly, partial correlation was used to assess the association between protein markers and the CSF NEFL level while controlling for the CSF HIV-1 level. Gene signature set enrichment analysis (GSEA) was used to calculate the enrichment score of the immune related pathways [149].

The immune related pathways were defined as the child terms of the term “immune system process GO:0002376” [150]. Because the **Olink Explore 1536** platform only measured a selected set of the human proteins, we only included an immune pathway if more than five of its proteins were measured by Olink assay. All statistical tests were adjusted for multiple testing using the FDR method.

Comparisons of the quadruplicate results of three proteins that were analyzed on each of the four plates and of earlier NfL measurements to the Olink NEFL results by linear regression used Prism 9.5.1 and 10.1, GraphPad Software, San Diego, California USA (www.graphpad.com). Selected comparisons of groups by ANOVA with correction for multiple comparisons and graphs of subject profiles were also produced using Prism 9.5.1 and 10.1.

## ACKNOWLEDGEMENTS

This study was supported principally by NIH/NINDS research grant 3R01 NS094067-05S and the Swedish state under an agreement between the Swedish government and the county councils (ALF agreement ALFGBG-965885). Previous research grants supporting CSF specimen collections and background data assays included: NIH/NIMH grant P01 MH094177, P01 DA026134, R01 MH62701, R01 MH081772, R01 NS37660, R01 NS43103, P01 DA026134, R01 NS37660, R01 DA051890, R01 MH62701, R01 MH096619, RR18522,nd P30 AI027763 and NCRR UCSF-CTSI UL1 RR024131); Sahlgrenska Academy at University of Gothenburg (project ALFGBG-11067); and the Swedish Research Council (project 2007-7092.3). HZ is a Wallenberg Scholar and a Distinguished Professor at the Swedish Research Council supported by grants from the Swedish Research Council (#2023-00356; #2022-01018 and #2019-02397), the European Union’s Horizon Europe research and innovation programme under grant agreement No 101053962, Swedish State Support for Clinical Research (#ALFGBG-71320), the Alzheimer Drug Discovery Foundation (ADDF), USA (#201809-2016862), the AD Strategic Fund and the Alzheimer’s Association (#ADSF-21-831376-C, #ADSF-21-831381-C, #ADSF-21-831377-C, and #ADSF-24-1284328-C), the Bluefield Project, Cure Alzheimer’s Fund, the Olav Thon Foundation, the Erling-Persson Family Foundation, Stiftelsen för Gamla Tjänarinnor, Hjärnfonden, Sweden (#FO2022-0270), the European Union’s Horizon 2020 research and innovation programme under the Marie Skłodowska-Curie grant agreement No 860197 (MIRIADE), the European Union Joint Programme – Neurodegenerative Disease Research (JPND2021-00694), the National Institute for Health and Care Research University College London Hospitals Biomedical Research Centre, and the UK Dementia Research Institute at UCL (UKDRI-1003).

## Supplemental Figure

**Figure S1.**
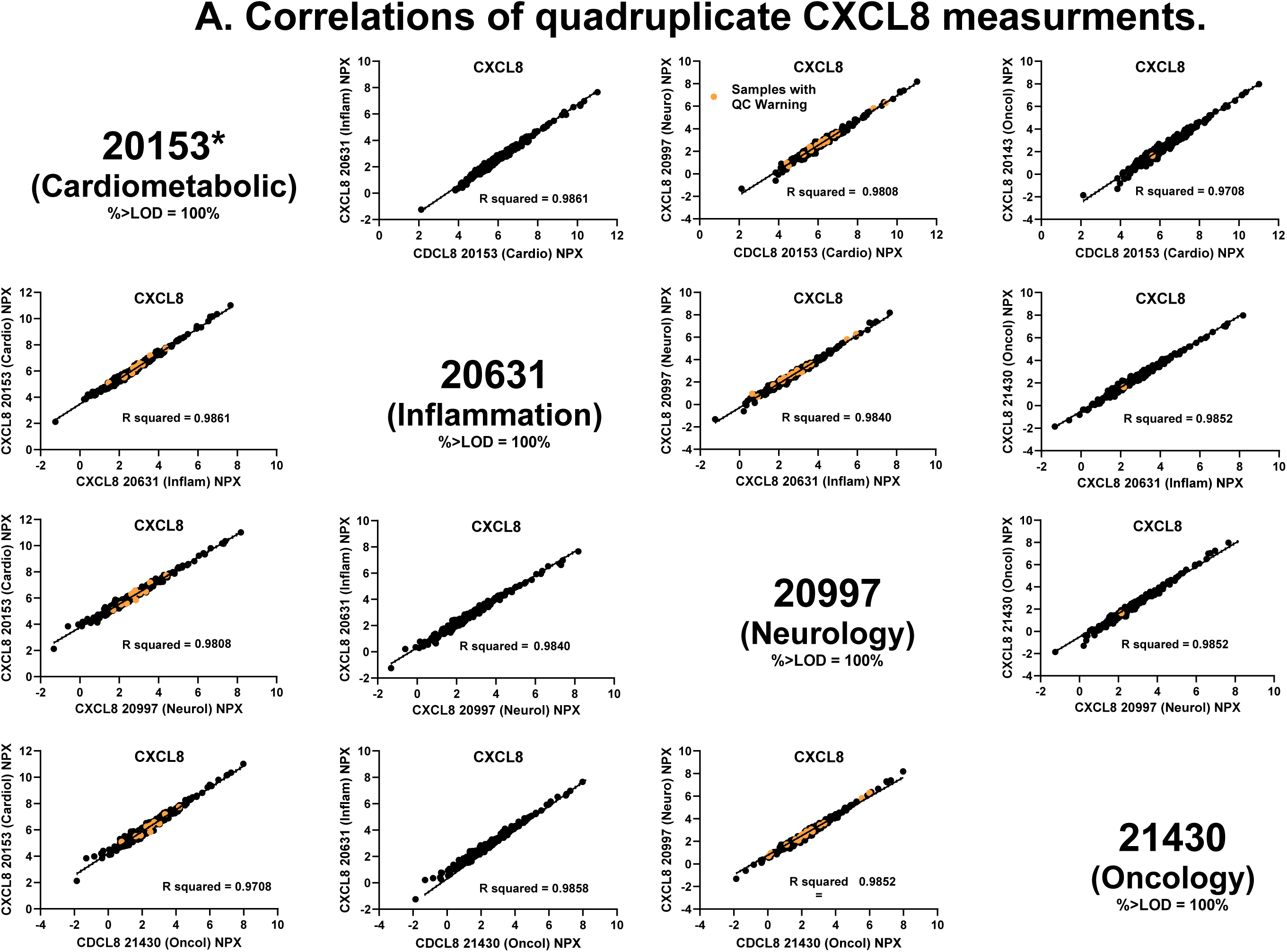
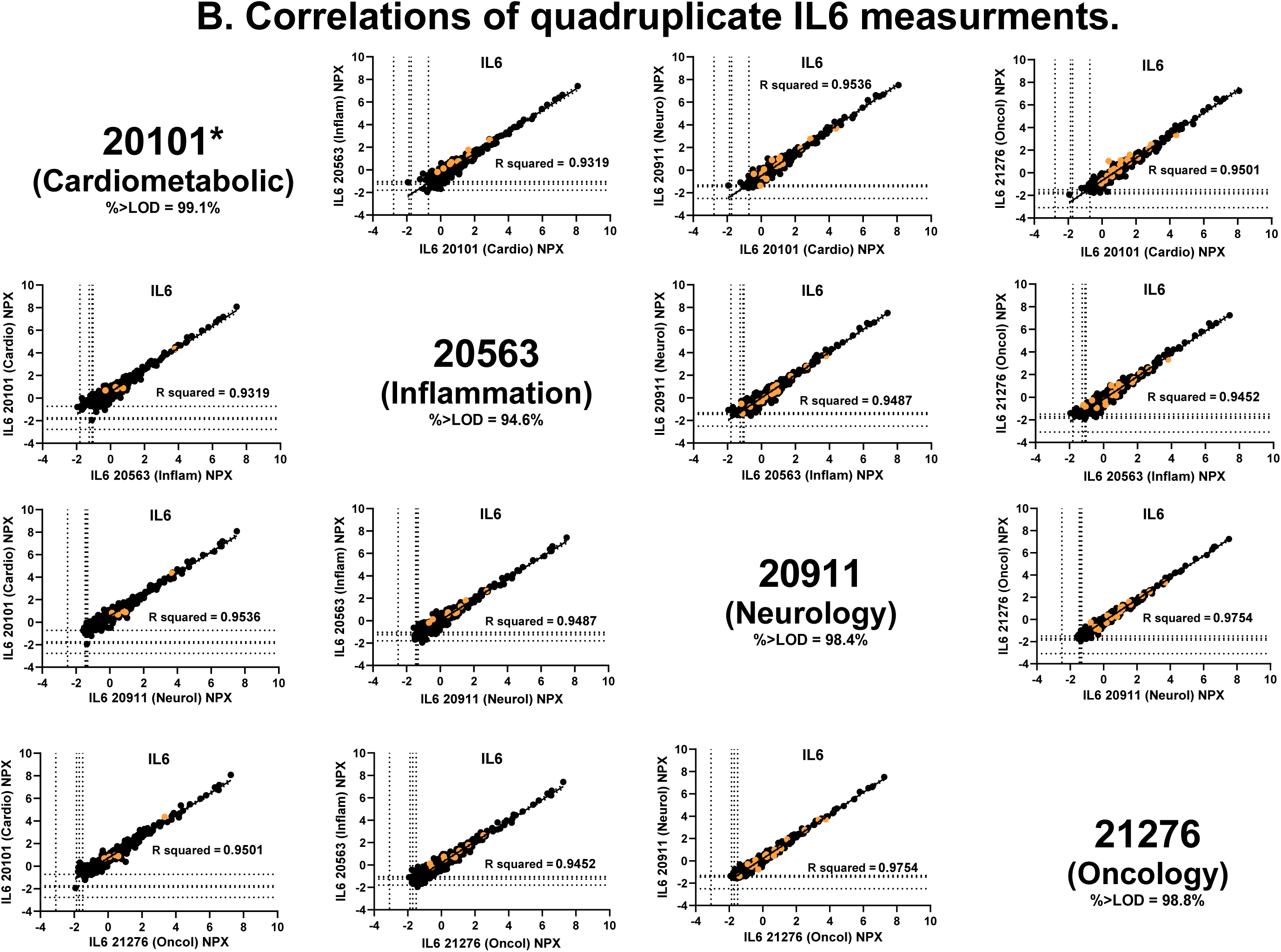
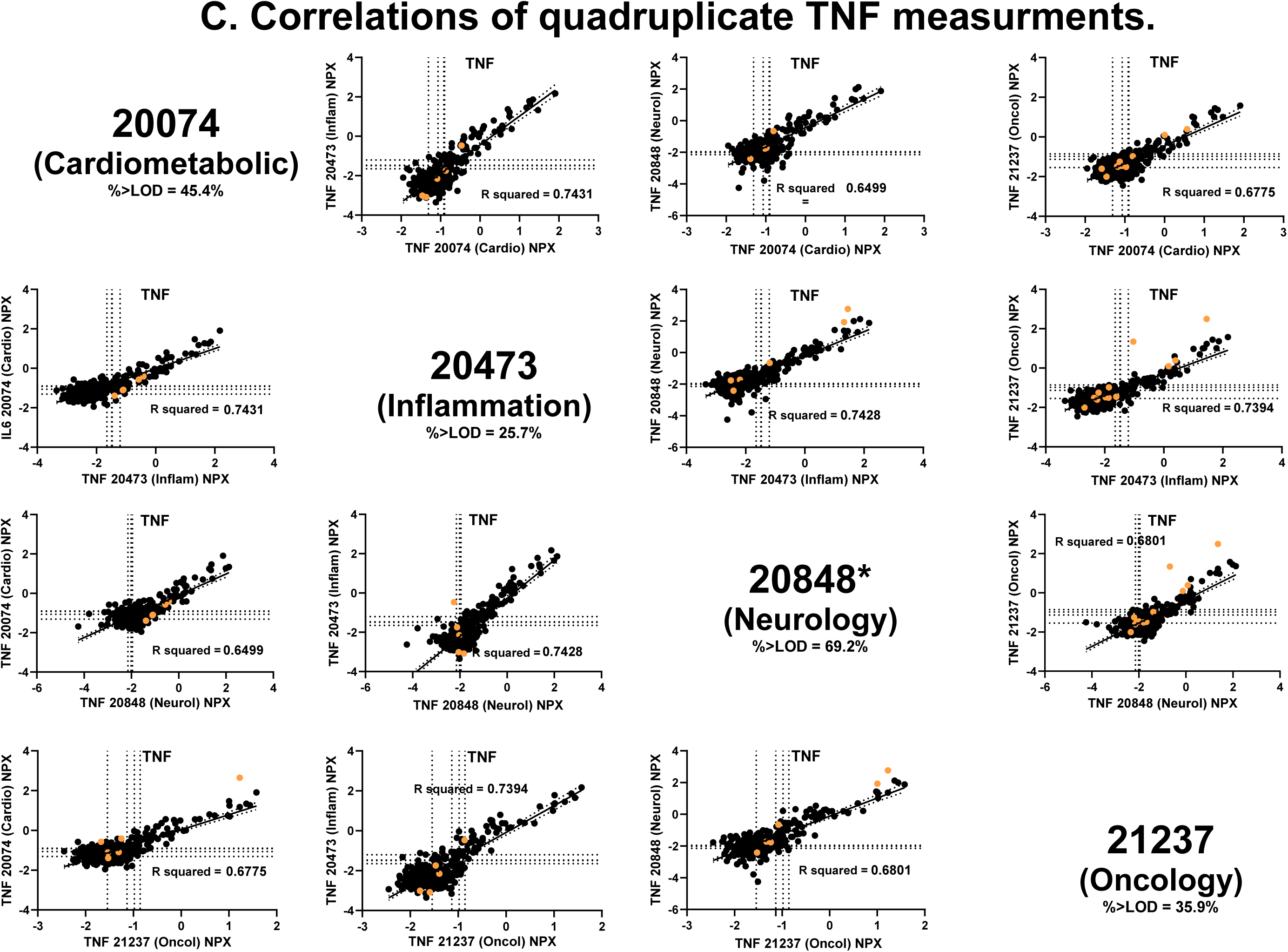
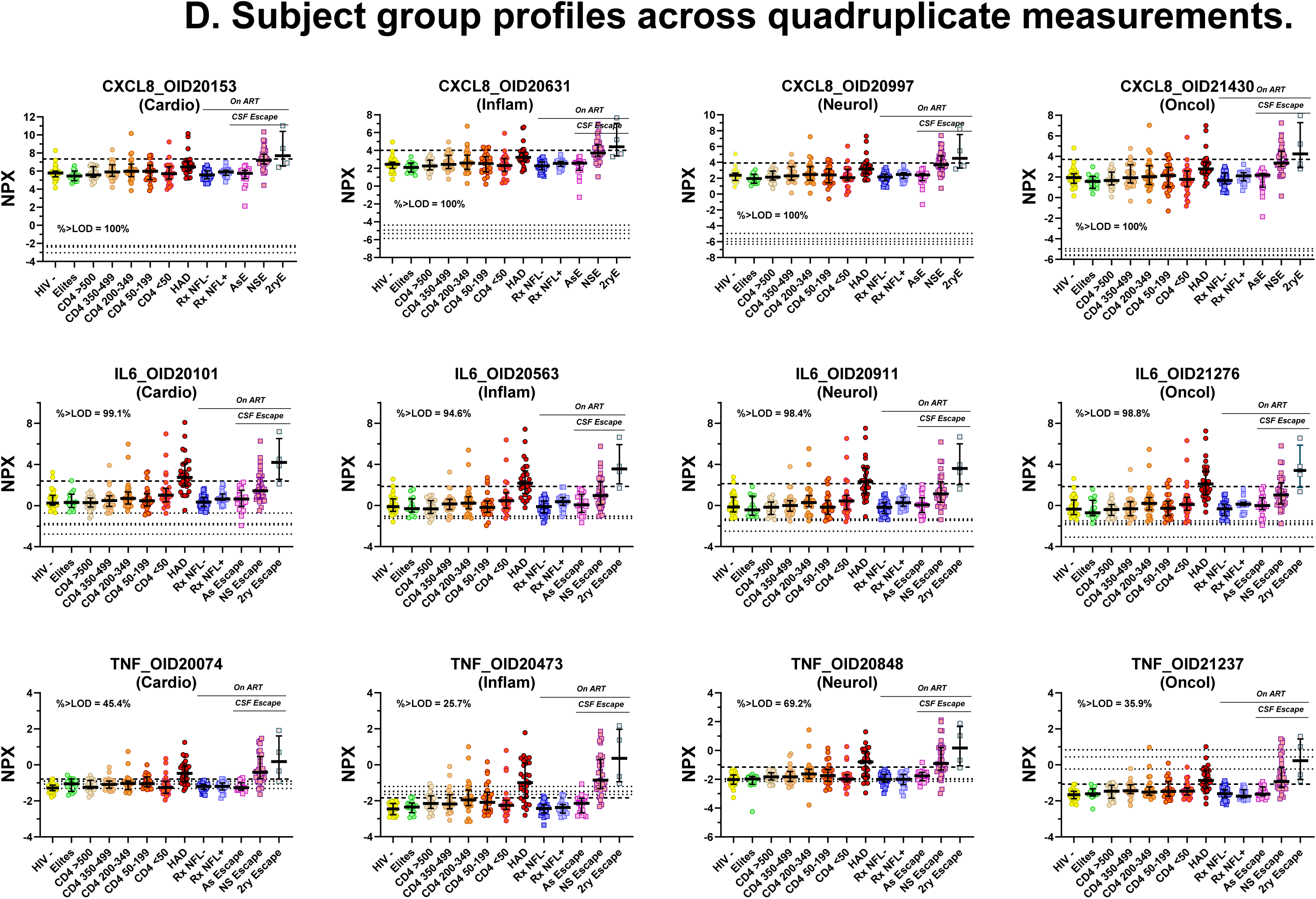
Quadruplicate measurements of CXCL8, IL6 and TNF. The **Olink Explore 1536** platform used four individual assay plates (*Cardiometabolic*, *Inflammation*, *Neurology* and *Oncology*), each with different sets of protein detection antibodies (two antibodies for each protein measured). As part of the quality control strategy of this platform, three of the proteins (CXCL8, IL6 and TNF) were independently measured on all four plates, i.e., in quadruplicate, using the same antibody pairs, but with each plate containing a different background of other antibody pairs. This supplemental figure examines the intercorrelations among these assay results and compares the patterns of study specimen profiles across the subject groups for each of these quadruplicate assays. This provides a view of the correlations among the measurements, the (negligible) impact of assays with QC Warnings, and the effect of some of the values below the LODs for particular measurements. ***Panels S1 A-C***. **Correlations among quadruplicate plate assay results**. This series of figures includes three montages, each with 12 graphs showing assay correlations among the quadruplicate assays along with identification of the QC Warnings related to individual measurements. The format for the first set (CXCL8) is outlined in detail that can then be applied to the other two sets of quadruplicate assays (IL6 and TNF). ***S1A***. ***Correlation of CSF CXCL8 measurements in quadruplicate assays***. This panel shows the correlations of the results of four assays (one from each plate) of the CSF CXCL8 concentrations in NPX units. The assay result numbers and plate names in the four spaces without graphs refer to the protein results graphed on the X axis of the panels in that row, and also on the Y axis of the panels in the column, e.g., measurement 20153 was obtained on the Cardiometabolic plate and these results were used in the X axis of the other graphs in the (top) row and the Y axis of the graphs in the (left) column. The panels show both the close correlations of the results across the quadruplicate measurement and the variability of absolute results in NPX units among the plates, and, indeed, differences in the scales of results among plates related to the effects of the differences in the background antibodies in each of the plates. The graphing of the points with QC warnings shown (orange symbols) relate to the measurement shown in the column of the panel and to the protein on the Y axis (for example, the QC Warnings for the Cardiometabolic plate warnings for its CXCL8 assay were included in the three graphs below this label). With respect to the correlations, there were only 6 sets of correlations, but the other 6 panels show graphs of the same variable pairs mirrored across the diagonal of the panel names. These graphs show the QC Warnings of different assays (those of the columns). The six linear correlations performed across these results were all very tight with an overall mean R^2^ = 0.9821 +/- SD 0.0059. The “**\***” on the upper left box (20153 assay of CXCL8 on the Cardiometabolic plate) indicates that the results for the CXCL8 measurements from this plate were used in the analyses presented in this report (and highlighted by gold-yellow background in **Table S1**). All of the QC Warning samples were within the body of other assay results. Since, as shown in the non-graph panels, all results were above the LODs for that measurement, the LODs are not shown in this plate. ***S1B***. ***Correlation of CSF IL6 measurements in quadruplicate assays***. This panel showing the correlations among the four CSF IL6 assays (***Figure S1B***) is arranged in the same format as the CXCL8 panels discussed above and uses the same labelling rules. Again, the correlations among the 6 assays were high with a mean R^2^ mean of 0.9508 +/- SD 0.0142. The measurements with QC warnings were again all well within the body of the overall data. For IL6, a small portion of the measurements were below the assay LODs (the percent for each of the assays is listed in the non-graph panels). The four LODs for each assay are shown with dotted horizontal and vertical lines. While the figure does not allow distinction of which data points relate to to the the individual four LODs, in general only a relatively small number of low values were near or below the LOD values, and they therefore likely had only a very small effect on the listed correlations. ***S1C***. ***Correlation of CSF TNF measurements in quadruplicate assays***. This panel (***Figure S1C***) is arranged in the same format as the CXCL8 and IL6 results discussed above. However, in contrast to those other results, the TNF intercorrelations among the quadruplicates were distinctly lower, and the mean R^2^ was 0.7055 SD +/- 0.0412. Additionally, two of the measurements with QC Warnings were outliers; this is most easily visualized in the two graphs in the Oncology plate results column (middle two subpanels on the right). More importantly, however, many more of the results were below their LODs, most notably in the Oncology plate (lower row) with only 35.9% of measurements above their related LODs. This, at least partially, contributed to the lower intercorrelations among the four TNF measurements. ***S1D***. **Subject group profiles of the three sets of quadruplicate CSF protein measurements across the study subject groups**. This group of graphs displays the results of each of the 3 quadruplicate assays (CXCL8, IL6 and TNF in the top, middle and bottom rows of graphs, respectively) for all of the subjects in this analysis. They are derived from the same measurements as the above panels but displayed according to the subject groups using the same format (including color coding of symbols and median +/- SEM error bars) for other group profiles in this report as outlined in Figure 1. In each of these panels the horizontal dashed line is set at the mean + 2SD of the HIV-controls for visual reference across the 13 groups while the four dotted lines show the four LODs for each of the assays which was most striking with the Oncol plate (lower right). The patterns of CSF protein changes across the groups were very similar for each of the quadruplicate measurements of the individual proteins (CXCL8, IL6 and TNF), including in relation to the HIV-control values, changes in medians across the CD4 defined groups as blood CD4 cells fell, elevations of HAD and NSE relative to each other (for example: modest elevations of CXCL8 with median values of ***NSE*** > ***HAD***; elevations of IL6 in ***HAD*** with relatively low values for **NSE**; and nearly equal elevations of both ***HAD*** and ***NSE*** for TNF). While for each of the four replicates the relative changes in the subject groups were largely the same, there were notable differences across some of the assays in the absolute NPX values and ranges. For example, the NPX ranges and values in the four panels of CXCL8, particularly between the right and left panels (Cardio and Oncol results), differed. This was less marked for the IL6 repeats, but there were notable differences in the HIV-negative medians in the left two panels of TNF results (Cardiology and Inflammation). This may have obscured distinctions among the ***CD4-defined*** and aviremic groups for this cytokine. These comparisons thus clearly illustrate that the NPX measurements are *relative* measurements *within each assay* that differed among the quadruplicates, yet showed similar relative changes with respect to group comparisons. They also show the potential attenuating impact of measurements below the LODs for the assays. **Glossary of gene-protein functions of the quadruplicate proteins** (extracted from https://www.uniprot.org/uniprotkb). These brief excerpts outline some of the known functions of the listed proteins, though they may not directly apply to their actions in CSF in the HIV-1 infection setting. **CXCL8 (*P10145***). (*Interleukin-8*). This is a chemotactic cytokine that mediates inflammatory response by attracting neutrophils, basophils, and T-cells. It is released in response to inflammatory stimuli, exerts its effect by binding to the G-protein-coupled receptors CXCR1 and CXCR2, and is found in neutrophils, monocytes and endothelial cells. Changes in CXCL8 concentrations across the specimen groups were modest with a shallow *lymphoid* pattern across the ***CD4-defined*** groups, and higher concentrations in the ***NSE*** than ***HAD*** groups. **IL6**. (***P05231***) (*Interleukin-6*). This cytokine has a wide variety of biological functions in the innate immune response, is synthesized by myeloid cells, including macrophages and dendritic cells, upon recognition of pathogens through toll-like receptors at the site of infection or tissue injury. In the adaptive immune response, it is required for the differentiation of B cells into immunoglobulin-secreting cells. It plays a major role in the differentiation of CD4^+^ T cell subsets. IL6 showed a rather flat pattern across the ***CD4-defined*** groups, modest elevation in the ***HAD*** group that was higher than the small increase in the ***NSE*** group. Highest medians were in the small ***2ryE*** group. **TNF**. (***P01375***) (*Tumor necrosis factor, also TNF-alpha, cachectin*). TNF is a multifunctional proinflammatory cytokine, mainly secreted by macrophages and involved in the regulation of a wide spectrum of biological processes. Here also the changes across the ***CD4-defined*** groups were shallow, perhaps suggesting a low-level *lymphoid* pattern, though also possibly attenuated by the frequency of values below the LODs. The ***HAD*** and ***NSE*** showed nearly equal mild, though distinct, elevations. Highest median was in the small ***2ryE*** group.

**Figure S2.**
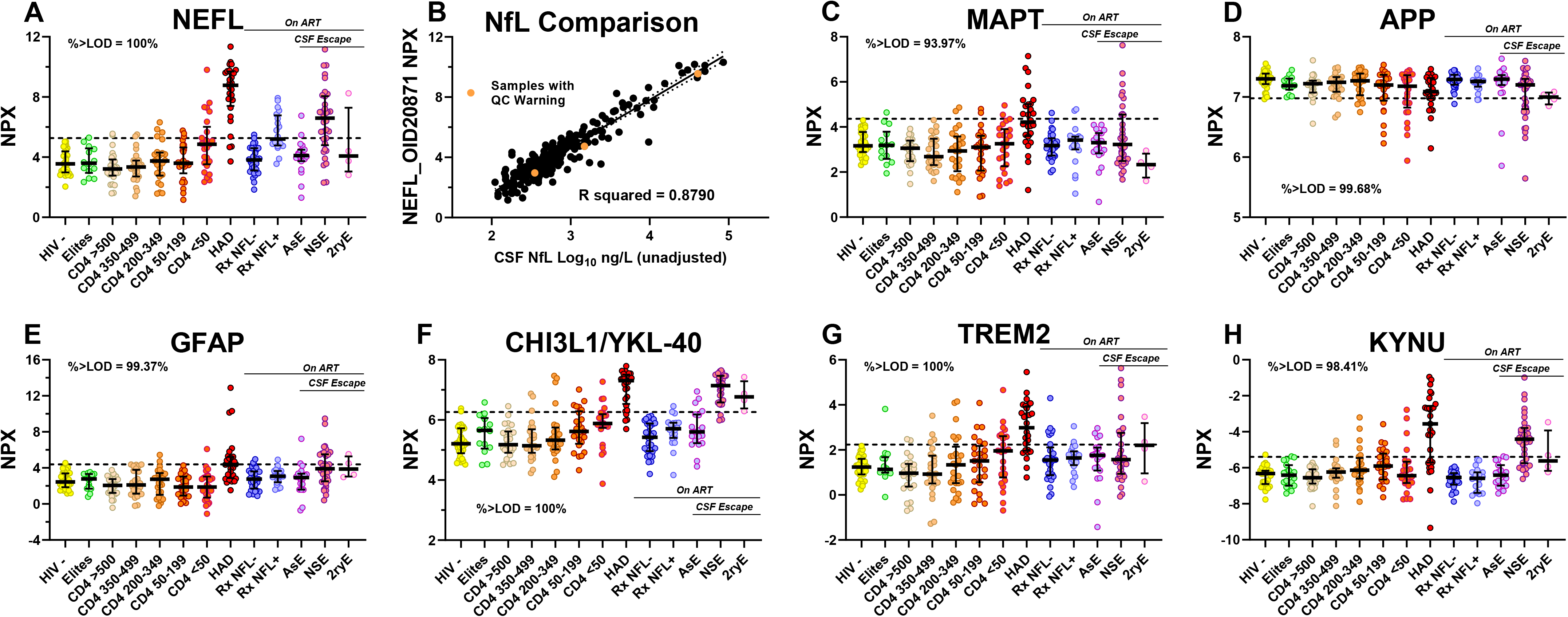
CSF protein biomarkers of CNS injury measured in Olink Explore platform. The figure shows the subject group profiles for seven **Olink Explore 1536** protein measurements that have been applied to assess CNS injury in other studies. In **Panels A** and **B – H**, the format is the same as that for other CSF biomarker changes across the groups, including the horizontal dashed line that indicates the mean + 2 SD of the ***HIV-*** controls as a visual reference for comparison across group (or – 2 SD in the case of APP in **Panel D** in which the concentration changes in the PWH were negative). Following the nomenclature used in the **Olink Explore 1536** platform, the proteins are designated by their gene names. This includes the Olink measurement of NfL which in this paper we refer to as *NEFL* using the gene designation as with the other assays. This distinguishes the Olink assay from the results of the UMAN ELISA used for the earlier measurements that were presented in Figure 1, **Panel O** for which we use the more general term abbreviation, *NfL*. **A. NEFL** (P07196) (*Neurofibrillary light chain protein*, also *NfL*). This is a major intracellular axonal protein that is shed into the CSF after neuronal-axonal injury. It has proved very useful as a CSF (and blood) biomarker of CNS injury in a number of neurodegenerative conditions [65] including HIV-1 infection [68]. This panel shows that the pattern of concentration changes of NEFL measurements generated in the **Olink Explore 1536** were very similar to those of the UMAN ELISA results presented earlier in Figure 1, **Panel O**. Greatest elevations were present in the ***HAD*** group, minor increases in the ***CD5 <50*** group, and substantial elevations in the ***NSE*** group (though lower than in ***HAD***). The ***Rx NFL+*** group showed clear elevation with a median near the level of the mean + 2 SD dotted line. This pattern of change might be considered as the prototypic *neuronal* pattern of change across the sample group. It is similar to the *myeloid* pattern, though perhaps distinguished by the flat concentration levels in the ***CD4-defined groups*** with CD4+ T cells above 50 per µL, along with elevation in the ***NSE*** but not the ***2ryE*** group. Whether the general elevation in the ***Rx NfL+*** group should be considered a part of this pattern or is peculiar to the NfL assay results requires further understanding of this group. These minor distinctions from the *myeloid* pattern also require further analysis and validation. **B. Correlation of CSF NEFL (Olink) with prior NfL measurements without age adjustment.** The regression shows the high correlation between the Olink NEFL and the prior Uman NfL ELISA assay results in the subset of the subjects presented in Figure 1, **Panel O** but without age adjustment. In this comparison high correlation was noted (R^2^ = 0.8790) despite the earlier measurements having been performed in multiple laboratory runs over several years. As in **Figure S1**, comparing analysis of the quadruplicate repeated assays, the NEFL measurements with QC Warnings, shown by orange symbols, fell within the central body of the results along the regression line, further supporting the inclusion of measurements with QC Warnings in our analysis. **C. MAPT (P10636**). (*Microtubule-associated protein*, also *tau*). CSF levels of MAPT showed a more blunted response to HIV-related injury than NEFL, as previously reported [123, 154, 155]. MAPT promotes microtubule assembly and stability in neurons and participates in the establishment and maintenance of neuronal polarity. It forms the neurofibrillary tangles characteristic of Alzheimer’s Disease. In this figure, only the ***HAD*** group exhibited a clear median increase with about half of the results above the 2 SD level of the control mean. The median of the ***NSE*** group was similar to the control group with a tail of about 25% of the samples falling above the 95% confidence limit. Hence, this protein appeared to be a less sensitive marker of neural injury than NEFL in HIV-1 infection. **D. APP** (P05067). (*Amyloid-beta precursor protein*). APP functions as a cell surface receptor and performs physiological functions on the surface of neurons relevant to neurite growth, neuronal adhesion and axonogenesis. Interaction between APP molecules on neighboring cells promotes synaptogenesis. Measurement of this protein in the panel showed only a modest decrease in median level in the ***HAD*** group, as previously reported [155] and a seeming lower median in the ***2ryE*** group. Thus, both MAPT and APP CSF measurements, which serve as indicators of Alzheimer’s disease, are not sensitive to HIV-1-related CNS injury, likely because of fundamental differences in their neuronal pathologies. **E. GFAP (*P14136***). (*Glial fibrillary acidic protein*). GFAP is a cell-specific intermediate filament that distinguishes astrocytes from other CNS cells. CSF GFAP concentrations were normal in most of the groups except for the ***HAD*** group and to a lesser extent the ***NSE*** group with about 50% and <50% concentrations above the controls +2 SD line, respectively. Either GFAP was a relatively insensitive biomarker of astrocytic reactions in this setting, or astrocytic changes were a less prominent aspect of HIV-induced neuropathology than neuronal injury (as documented by CSF NEFL changes). **F. CHI3L1/YKL-40** (***P36222***). (*Chitinase-3-like protein 1*). Systemically CHI3L1/YKL-40 plays a role in T-helper cell type 2 (Th2) inflammatory response and IL-13-induced inflammation, inflammatory cell apoptosis, dendritic cell accumulation and M2 macrophage differentiation. It has also been implicated as an astrocytic biomarker in CSF. CHI3L1/YKL-40 concentrations in CSF were distinctly elevated in *HAD* and *NSE* with an overall pattern similar to that of NEFL, but with an earlier median increase in the *CD4-defined* groups. Incidentally, the value in the ***Rx NFL+*** group was similar to that of the ***Rx NFL-***, which might imply that if injury was present in the ***Rx NFL+*** group, it was circumscribed, affecting axons but sparing other cellular elements. The small ***2ryE*** group values were also elevated, presumably related to strong glial reaction in these individuals with a *myeloid* pattern in the ***CD4-defined* groups**. Overall, CHI3L1/YKL-4 appeared to be a sensitive biomarker of HIV-1-induced pathology and perhaps a useful companion to NfL. **G. TREM2** (***Q9NZC2***). (*Triggering receptor expressed on myeloid cells*). TREM1 acts as a receptor for amyloid-beta protein 42, a cleavage product of the amyloid-beta precursor protein APP, and mediates its uptake and degradation by microglia. In this study, there was an increase in TREM2 mean values in the lower ***CD4-define***d groups and highest levels in the **HAD** group, but largely normal levels in ***NSE*** except for some elevated outliers possibly indicating a difference in their pathological substrates. This is consistent with TREM2’s putative association with microglial activation and the myeloid predominance in ***HAD*** but not in ***NSE***. The gradual increase in its concentration with falling CD4+ T cells, including the ***CD4 <50*** group, also shows the *myeloid* pattern. This differences from CHI3L1/YKL-4, perhaps most notably the absence of elevation in the NSE group suggests that these two markers detect different cell changes, with YKL-40 importantly encompassing astroglial changes present in both ***HAD*** and ***NSE***, while TREM2 largely may lindicate, myeloid activation that is prominent in ***HAD*** but not in ***NSE***. **H. KYNU** (***Q16719***). (*Kynureninase*). This is an enzyme in L-kynurenine pathway considered to contribute to inflammation and neuronal injury in ***HIVE***. There was a prominent increase in ***HAD*** and lesser increase in ***NSE*** with little change from control in the other groups. The pattern in ***CD4-defined*** groups was perhaps ambiguous, but the overall median values of these groups were close to that of the HIV-controls. KYNY could possibly be useful as a marker of severe CNS injury in HIV-1 infection. The following table lists the correlation of NEFL with the other Olink measurements.

---

| Protein<br>(UniProt) | Other Names | Correlation w NEFL |  | Description |
| --- | --- | --- | --- | --- |
|  |  | P value | R<sup>2</sup> |  |
| **NEFL** | Neurofilament light chain prot | - | - | Axonal neurofilament protein |
| **MAPT** | Tau | <0.0001 | 0.3471 | Binds axonal microtubules |
| **APP** | Amyloid-β precursor protein | 0.1946 | 0.0058 | Neuronal cell surface protein |
| **GFAP** | Glial fibrillary acidic protein | <0.0001 | 0.4545 | Astrocyte intermediate filament |
| **CHI3L1** | Chitinase-3-like prot 1, YKL-40 | <0.0001 | 0.6005 | Astroglial and microglial cell |
| **TREM2** | Triggering receptor myeloid cells 2 | <0.0001 | 0.5622 | Myeloid cell (microglial and MΦ) activation |
| **KYNU** | Kynureninase | <0.0001 | 0.3690 | Tryptophan metabolism, KYN & QUIN |

**Table S1 Notes**

This Table contains the full study data set and these notes are intended to serve as a brief guide to the reader and, more particularly, as an aid to other investigators who use these data for additional analysis. The Table is divided into two main sections: the first (***columns A – AD***) presents the salient *background features* of the study participants, including demographic features, HIV-1-related laboratory findings and routine CSF laboratory results. The larger remaining section (***columns AE – BEV***) lists the **Olink Explore 1536** *protein measurements* in NPX units.

**Background features of study groups and specimens.**

The CSF samples were segregated into clinically- and biologically-defined groups (**A** – **N**) that included: HIV-seronegative controls (**A**); 6 HIV-1-infected, untreated (**B – H**); and 5 HIV-1-infected, ART-treated (**I – M**) main groups. CSF proteins from a miscellaneous group of four individuals (**N**) who did not fit any of the main categories, were also measured but their results were *<u>not included in this analysis</u>* because of its focus on group identities and differences. The background definitions of the study groups are outlined in the **Methods** section and their features further described in the **Results-Discussion** section of the manuscript. Within these groups, individual study subjects are then numbered. In addition to this Group-Subject Code identifier, earlier <u>Subject IDs</u> and <u>Sample IDs</u> are also included in the Table for cross-referencing to use of related specimens in other studies. The untreated PLWH are classified on the basis of virological status (***elite*** controllers with plasma HIV-1 RNA below detection), CD4 counts as a measure of systemic disease progression in neurologically asymptomatic individuals (divided into five ***CD4-defined*** brackets), and a group identified by abnormal neurological status (***HAD*** patients who were studied during their clinical disease presentation). The ART-treated PLWH included two groups with plasma viral suppression (<20 copies of HIV-1 RNA): a more representative group with CSF NfL measurements that were either normal on earlier measurement or not previously assessed (***RxNFL-***), and a more unusual selected group identified by CSF NfL elevation on prior testing (***RxNFL+***). The remaining three treated groups were selected because of HIV-1 CSF escape categorized into three clinical types: asymptomatic, neurosymptomatic and secondary (***ASE***, ***NSE*** and ***2ryE***).

The left-sided section of the Table (columns **A** – **AD**) lists the basic demographic, HIV-1-related laboratory (e.g, blood and CSF HIV-1 RNA measurements, blood T cell counts), and clinical CSF (WBC counts, protein concentrations, CSF:blood albumin ratio) information from these individuals. Additionally, for some CSF and blood neopterin and CSF NfL measurements that were available from earlier study.

The abbreviations of drugs and combinations listed in column K follow usage outlined in Key to Acronyms in*Guidelines for the Use of Antiretroviral Agents in Adults and Adolescents with HIV*, https://clinicalinfo.hiv.gov/en/guidelines/hiv-clinical-guidelines-adult-and-adolescent-arv/drug-name-abbreviations, updated December 06, 2023.

**Olink Explore output.**

The remainder of the Table (columns **AE** – **BEV**) presents the individual Olink CSF protein measurements in NPX units. Proteins are listed in alphabetical order and, following the Olink usage, are identified by the gene nomenclature and number IDs as listed in UniProt (https://www.uniprot.org/uniprotkb/).Where the Olink data report indicated a *Quality Control (QC) Warning*, the individual protein data cells are highlighted in light tan. The data with QC Warnings are all included in the analysis presented in this paper.

The column titles for the three quadruplicate protein assays (CXCL8. IL6 and TNF) are highlighted in yellow, with the individual assay results chosen at random for use in this analysis highlighted in a darker golden yellow; the other three assay results for each of these three proteins are omitted from the analysis except as described in the exploration of their intercorrelation and effect of QC Warnings (see **Figure S1**).

As noted in the two columns before the Olink results (columns **AE** and **AF**), the Table also annotates the LODs for each assay and their potential impact on the assay results. Each protein assay was comprised of four sections, each with its own LOD. In this Table, these are listed in rows **325** – **328** within the column for measured protein. To allow the user to align each protein assay with its related LOD, the LODs have been labelled ***A***, ***B***, ***C*** or ***D***, and each protein measurement has similarly been labelled with the pertinent group letter in this column. As an overall index of the relationship of the individual protein measurement to the related LODs, the percentages of values above the related LOD are listed in row **329** for each protein. These are also color-coded in 6 brackets: **100%**; **90% - <100%**; **60% - <90%**; **40% - <60%**; **20% - <40%** and **<20%**.

Omitting the four miscellaneous subjects reduced the data included in the current analysis to a total of 303 unique CSF samples segregated into 13 clinically-defined groups. while omitting three of each of the three quadruplicate measurements reduced the Olink output to measuring 1,463 unique proteins. This resulted in a final total of 443,289 Olink protein measurements contributing to the analysis.

**Table S2.**
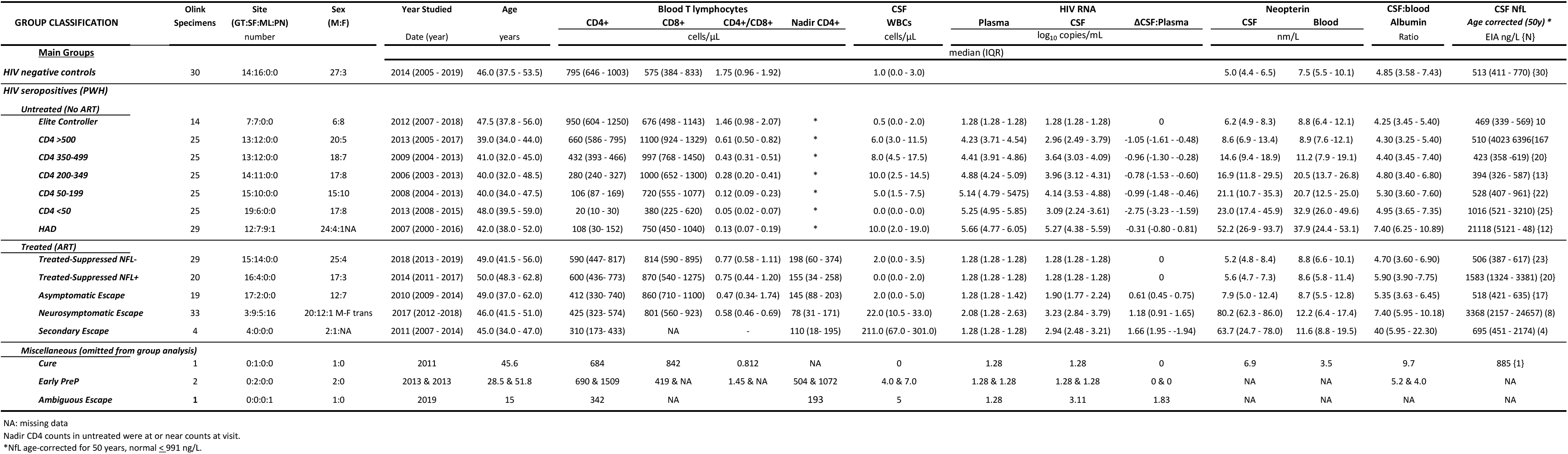
Background characteristics of the study groups.

